# Stepwise recruitment of Hsc70 by DNAJB1 produces ordered arrays primed for bursts of amyloid fibre disassembly

**DOI:** 10.1101/2024.01.25.577078

**Authors:** Jim Monistrol, Joseph G Beton, Erin C Johnston, Helen R Saibil

## Abstract

To understand the action of co-chaperones of the J-domain protein family in the assembly of disaggregation-active Hsc70 complexes on the surface of amyloid fibres, we used cryo-EM and tomography to compare the assemblies with wild-type co-chaperones or an activity deficient mutant. We show that human DNAJB1 binds uniformly and densely along α-synuclein amyloid fibres in an asymmetric orientation, with one subunit of the DNAJB1 dimer lying along the fibre surface and the other subunit further away. It acts in a 2-step recruitment of Hsc70 to the fibres, first releasing DNAJB1 auto-inhibition and then activating the Hsc70 ATPase by the J domain, with ATPase recycling stimulated by the nucleotide exchange factor Apg2. When the auto-inhibition is removed by mutating the H5 inhibitory binding site on DNAJB1 (Δ1H5 DNAJB1 mutant), Hsc70 is recruited to the fibres at a normal level, but the resulting complex is inactive in disaggregation. Cryo tomography of the wild-type DNAJB1:Hsc70:Apg2:αSyn fibre complex shows dense arrays of the chaperones extending out from the fibre surface along spiral tracks, with the Hsc70 density on the outer surface of the DNAJB1 layer. The Δ1H5 DNAJB1:Hsc70:Apg2:αSyn fibre complex results in equally dense but less organised binding all over the fibre surface and lacks the ordered clusters. On the basis of these findings, we propose that 2-step activation of DNAJB1 regulates Hsc70 access to the fibre substrate. Since the DNAJB1 dimers are bound every 40 Å, the J domain released by Hsc70 binding to one dimer could activate the ATPase of an Hsc70 bound to the adjacent dimer. This could trigger a cascade of recruitment and activation in a localised, dense cloud of Hsc70 molecules to give coordinated, sequential binding and disaggregation from an exposed fibre end, as observed by cryo-EM and earlier fluorescence and AFM studies.

## Introduction

Neurodegenerative disorders are characterised by the accumulation of protein aggregates, for example those formed by α-synuclein (αSyn) in Parkinson’s disease (PD), dementia with Lewy Bodies and Multiple System Atrophy (Goedert et al., 2017). In these disorders, soluble αSyn monomers can self-assemble to generate unstable small species which can quickly react to form large more stable oligomers rich in β-strands and lastly highly ordered amyloid fibres (Cremades et al., 2012). Familial forms of PD, mainly caused by single point mutations or by duplication or triplication of the SNCA gene encoding αSyn, suggest a relationship between αSyn aggregation and neuronal death (Chartier-Harlin et al., 2004; Lesage et al., 2013; Polymeropoulos et al., 1997; Singleton et al., 2003). However, as in other protein aggregation diseases, the mechanism of αSyn toxicity remains unknown. Lewy bodies are cytosolic inclusions mainly composed of organelles, lipids and filaments (Shahmoradian et al., 2019). Fibrillar aggregates are present in post-mortem brains (Schweighauser et al., 2020) and both oligomers and fibrils are neurotoxic and can be transmitted between neurons, leading to disease propagation (Alam et al., 2019; Desplats et al., 2009; Peelaerts et al., 2015; Trinkaus et al., 2021). Moreover, amyloid fibres can promote αSyn aggregation by secondary nucleation (Iadanza et al., 2018).

The 70 kDa heat shock protein (Hsp70) family plays a critical role in the quality control system by regulating protein folding, activity and degradation, by preventing protein aggregation, and by acting in protein disaggregation (Nillegoda and Bukau, 2015; Rosenzweig et al., 2019). Hsp70 functions require cycles of association and dissociation with substrate via its ATPase activity. The Hsp70 ATPase cycle is regulated by two cochaperones: a J-domain protein (JDP) which defines the specificity of Hsp70 for the substrate and stimulates Hsp70 ATPase activity, and a nucleotide exchange factor (NEF) which controls ATPase cycle progression (Rosenzweig et al., 2019). In metazoa, the combination of Hsc70 (the constitutively expressed human cytosolic Hsp70), DNAJB1 (a class B JDP), and Apg2 (NEF Hsp110) can disassemble αSyn, tau and huntingtin exon 1 amyloid fibres by depolymerisation and fragmentation (Gao et al., 2015; Nachman et al., 2020; Scior et al., 2021). It is thought that the dense arrangement of Hsc70 chaperones bound at the fibril surface causes steric clashes between the chaperones and generates entropic pulling forces that destabilise the fibrils (Wentink et al., 2020). Fibrils are then disassembled unidirectionally by rapid bursts in which protofilaments are unwound and monomers are extracted (Beton et al., 2022; Franco et al., 2021; Schneider et al., 2021).

There are specific requirements for the cochaperones in amyloid disaggregation. Apg2 more efficiently enhances disaggregation than smaller NEFs such as Bag1 (Wentink et al., 2020); this is attributed to its larger mass. Apg2 also promotes the recruitment of dense clusters of Hsc70 at one end of the fibres (Beton et al., 2022). The specificity of DNAJB1, or at least class B JDPs, is also essential for disaggregation (Gao et al., 2015). The interactions between class B JDP and Hsc70 involve a 2-step mechanism (Faust et al., 2020). Initially, the C-terminal EEVD motif of Hsc70 binds DNAJB1. This triggers the release of the DNAJB1 auto-inhibitory helix V from the J-domain. The J-domain can then bind to the Hsc70 ATPase domain and stimulate ATP hydrolysis. Although the reason is still unclear, this 2-step mechanism is necessary for fibril disaggregation. In contrast, class A JDPs recruit Hsc70 in a 1-step mechanism that is ineffective in disaggregation of amyloid fibres (Faust et al., 2020).

The coordinated actions of these 3 chaperones can disaggregate amyloid fibrils *in vitro*. However, structural information about the mechanism is lacking. Here, we use cryo-electron microscopy (cryo-EM) to investigate the structure of DNAJB1 in complex with αSyn amyloid fibres. This reveals that DNAJB1 binds in a tightly packed array all along the fibres with a 40 Å repeat. This packing was also seen with a mutant of DNAJB1 lacking the J-domain and G/F linker (ΔJ-DNAJB1), which does not recruit Hsc70. A more targeted mutation replacing the autoinhibitory binding site on DNAJB1, making it functionally like a DNAJA (Faust et al, 2020) supports recruitment of Hsc70 to the fibres but not disaggregation, and does not yield the organised clusters of Hsc70 at fibre ends that appear to be essential for disaggregation.

## Results

### Low-salt conditions reversibly enhance DNAJB1 binding

DNAJB1 forms the initial interaction with αSyn amyloid fibres, binding densely all along the fibres (Gao et al., 2015). However, binding assays of 8 µM DNAJB1 to 20 µM αSyn amyloid fibres (monomer concentration) in physiological buffer show that only about 40% of the DNAJB1 interacts with the fibres (Fig 1A-B, condition in HKMD buffer). The remaining DNAJB1 appears as a high background of unbound protein, hampering image processing of the fibre-DNAJB1 complex. We therefore aimed to optimise DNAJB1 binding to the αSyn amyloid fibres. DNAJB1 binds to the acidic flexible C-terminus of αSyn and its interactions with the amyloid substrate are mostly electrostatic (Jiang et al., 2019; Wentink et al., 2020). Therefore, we reasoned that DNAJB1 binding could be increased by depleting salts from the buffer. We compared DNAJB1 binding in the high- and low-salt buffers (HKMD and HD, respectively). In HD buffer, using the same protein concentrations as above, 80% of the DNAJB1 binds to the amyloid fibres (Fig 1A-B), doubling the amount of DNAJB1 bound. Cryo-EM of the complex in HD buffer (Fig 1C) show a dense fuzzy decoration all along the fibres and a greatly reduced background, favouring this condition for single particle analysis. We checked that i) this low-salt buffer did not denature DNAJB1 by pre-incubating αSyn amyloid fibres with DNAJB1 in HD or HKMD buffer before performing a ThT assay in disaggregation buffer, including Hsc70 and Apg2 (Fig 1D) and ii) showed that the additional binding is reversed when the salts are added back to the buffer after a first incubation of αSyn amyloid fibres with DNAJB1 in HD buffer (condition HD/HKMD in Fig 1E-F). Since the HD buffer does not denature DNAJB1 and does not create irreversible interactions, this buffer was used to study the complex formed of WT DNAJB1 and αSyn amyloid fibres by cryo-EM.

**Figure 1.**
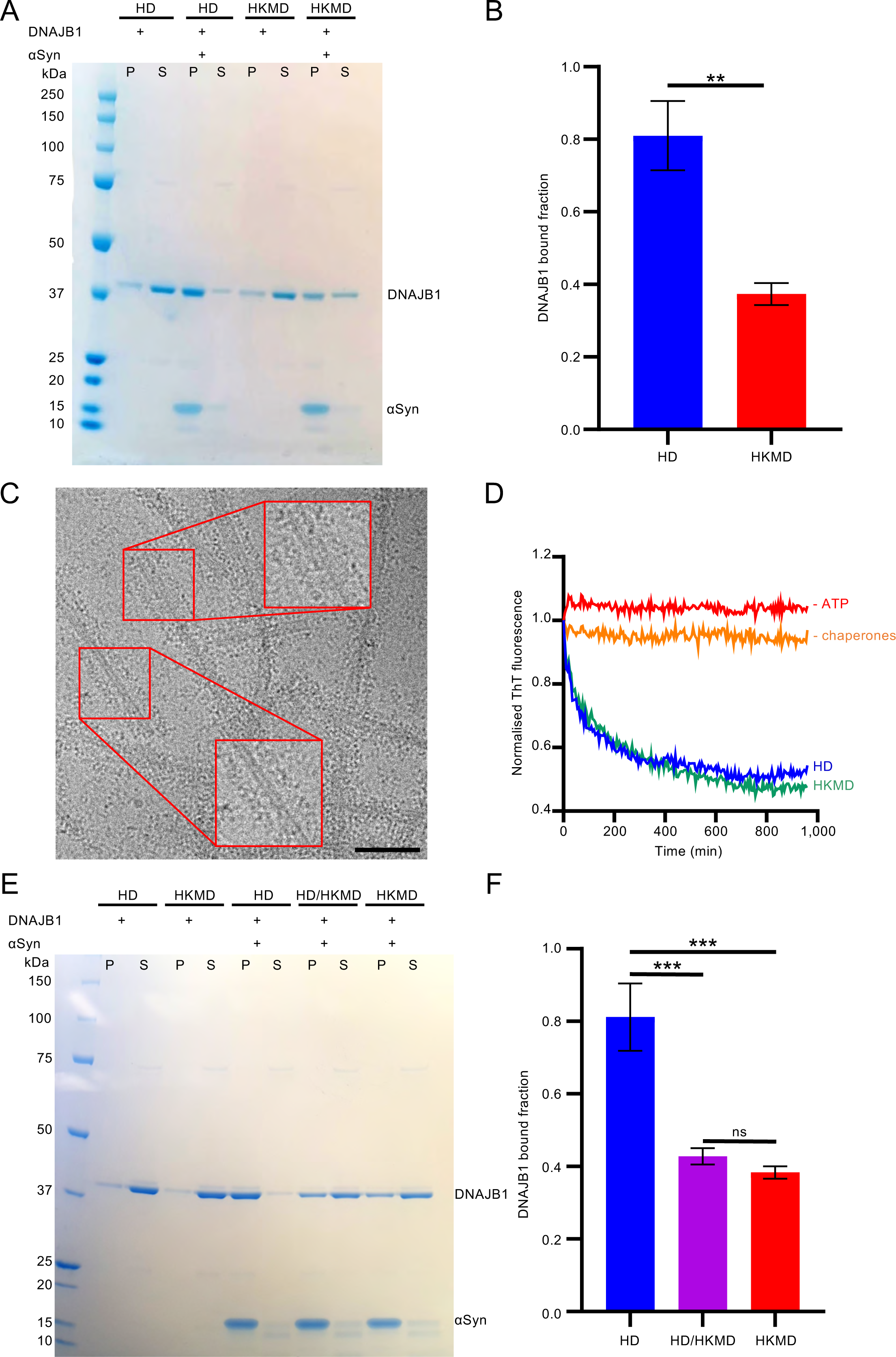
Effect of salt on DNAJB1 binding to αSyn amyloid fibres. A) Binding assay for WT DNAJB1 binding to αSyn amyloid fibres in HKMD or HD buffer. Two controls without amyloid fibres confirm that salt depletion does not cause precipitation of WT DNAJB1. A larger pellet band for DNAJB1 is observed in HD buffer in the presence of fibres. S: supernatant, P: pellet. B) Histogram of WT DNAJB1 bound fractions in each buffer (N = 3 independent experiments). A Shapiro-Wilk test was performed to check the normality of the data, followed by an unpaired t test (P = 0.0016). Mean DNAJB1 binding values with SD are shown. C) Cryo micrograph of αSyn amyloid fibres fully decorated by WT DNAJB1 in HD buffer. Scale bar, 50 nm. D) ThT assay showing that pre-incubation of WT DNAJB1 and αSyn amyloid fibres in HD buffer does not alter the disaggregation activity. The assay was performed in disaggregation buffer (table 1). E) Binding assay showing the reversibility of the binding when the salts are added back. The proteins were incubated twice in HD buffer (HD condition), twice in HKMD buffer (HKMD condition) or once in HD buffer and once in HKMD buffer (HD/HKMD buffer). F) Histogram of the WT DNAJB1 bound fraction in each condition shown in (E) (N = 3 independent experiments). A Shapiro-Wilk test was performed to check the normality of the data, followed by a one-way ANOVA with Tukey’s multiple comparisons test (P = 0.0004 between HD and HD/HKMD conditions, P = 0.0002 between HD and HKMD conditions). Mean Hsc70 binding values with SD are shown.

### DNAJB1 binds to αSyn amyloid fibres in an asymmetric position with a 40 Å repeat

We performed cryo-EM single particle analysis to study DNAJB1 binding to the fibres. The raw micrographs showed the fibres decorated with two layers of dots, with a ∼40 Å periodicity along the fibre axis (Fig 2A). This observation was confirmed on the 2D classes, in which the layered dots were more pronounced and clearly followed the helical twist of the fibres (Fig 2B-C). The corresponding diffraction patterns confirmed the 40 Å periodicity (Fig 2B). The resolution in the 2D class averages was low because the alignment was dominated by the amyloid fibre rather than the less ordered DNAJB1. Nevertheless, we were able to identify two fibre polymorphs within our dataset by combining visual inspection of 2D classes with the initial 3D model generation using the amyloid fibre reconstruction tool (relion_helix_inimodel2d) in RELION 3.1 (Supplementary fig 1). We separated the particles for each polymorph and processed them independently. Reconstruction of polymorph 1 showed clearly resolved DNAJB1 density. We generated a structure of this complex at ∼14 Å with a mask applied to focus on DNAJB1 decoration only on one side of the fibre (Fig 2D and Supplementary fig 2A). The DNAJB1 density shows a horseshoe shape oriented in an asymmetric position with one subunit closer to the fibre core. Fitting a model into this horseshoe density showed it is a plausible size and shape for DNAJB1 (Fig 2D) with the J-domains adjacent to the corresponding CTD (the autoinhibited form). A gap is visible between the fibril core and DNAJB1 peptide binding sites which are ∼60-90 Å away from the fibre surface. DNAJB1 interacts with the flexible C-terminus of αSyn, in particular residues 123-129 (Wentink et al., 2020). This leaves 30 disordered residues between the fibril core and the DNAJB1 binding site, which are sufficient to bridge the gap to the inner subunit of DNAJB1. Lastly, reprojections of the map were generated and compared with the 2D classes (Supplementary fig 2B-C). They show a reasonable resemblance, supporting the validity of the low-resolution map.

**Figure 2.**
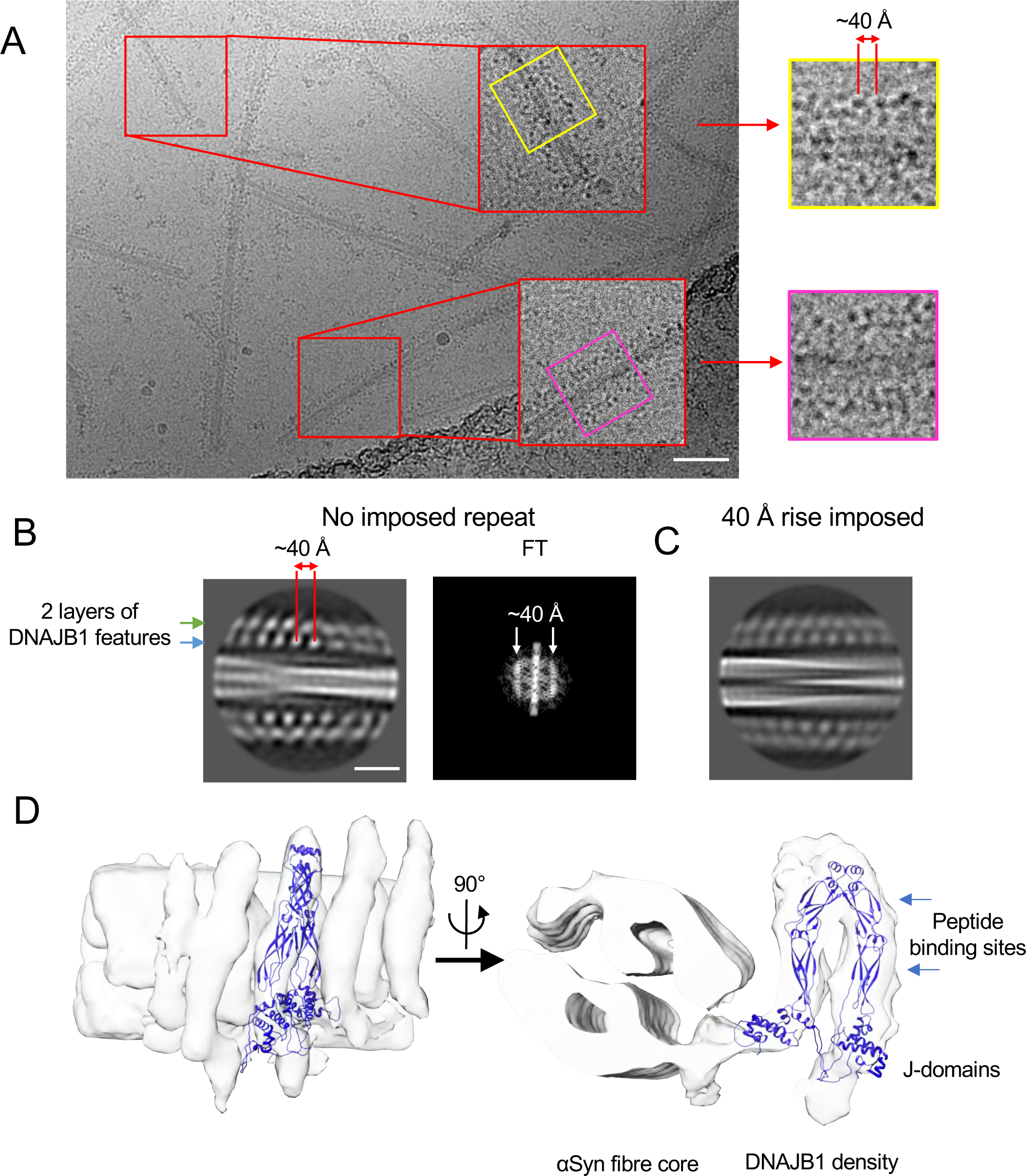
The complex of αSyn amyloid fibres and WT DNAJB1. A) Micrograph showing the fuzzy decoration of WT DNAJB1 on αSyn amyloid fibres. DNAJB1 repeats, displayed as dark dots, are separated by 40 Å. Scale bar, 500 Å. B) 2D class and FT showing the 40 Å repeat in WT DNAJB1 decoration, shown in reverse contrast. Scale bar, 100 Å. C) 2D class with a 40 Å repeat imposed for the helical rise. The 2D class is similar to the one displayed in (B) with 2 layers of dots characterising WT DNAJB1 decoration on the fibre. D) Reconstruction of the WT DNAJB1:αSyn fibre complex. The density corresponding to WT DNAJB1 was manually fitted with the crystal structure of the truncated human DNAJB1 lacking the J-domain and G/F linker (PDB: 3AGY) and the NMR structure of DNAJB1 J-domain (PDB: 1HDJ).

We also examined the αSyn amyloid fibres alone, by re-extracting the particles for each polymorph, without binning, and reprocessing. In this case, polymorph 2 gave clearer results. We classified the particles without imposing a subunit repeat and the corresponding 2D classes and diffraction pattern displayed the cross-β repeat of 4.8 Å typical of amyloid fibres (Supplementary fig 3A, B). We determined a 3D structure of this amyloid fibre at a nominal resolution of 3.6 Å (Supplementary fig 3C,D and Supplementary fig 4A,B). For polymorph 1, we could observe apparent side chain density in the core of the fibre, there was no visible cross-beta separation along the fibre axis. Nonetheless, we were able to identify that our reconstruction was similar to the previously reported polymorph 2b structure (PDB: 6SST, Guerrero-Ferreira et al., 2019)(Supplementary fig 1A and Supplementary fig 4C). The lack of cross-β separation in our reconstruction precluded model refinement, but we were able to rigid-body dock each protofilament (PDB ID: 6SST) with ChimeraX to generate an approximate model (Supplementary fig 4C).

### The J-domain is not required for DNAJB1 binding to the fibres

The asymmetric orientation of the JDP dimer on the fibre surface was unexpected. To see if the linker and J domain play a role in this binding geometry, we examined the complex formed by αSyn amyloid fibres with a truncated DNAJB1, lacking the G/F linker and J domain (ΔJ-DNAJB1). It has been shown that the DNAJB1 J-domain alone is sufficient to recruit Hsc70 to the fibres but does not support disaggregation (Wentink et al., 2020). We first assessed whether HD buffer would also promote ΔJ-DNAJB1 binding (Supplementary fig 5). A binding assay showed that 1) HD buffer doubles the amount of ΔJ-DNAJB1 bound to the fibres and 2) adding back the salts reverses the additional binding (Supplementary fig 5A,B). We also confirmed by cryo-EM that this condition promotes the formation of the complex with little ΔJ-DNAJB1 free in solution (Supplementary fig 5C). We did not assess the activity because it has already been shown that the J-domain is essential for recruitment of Hsc70 to αSyn amyloid fibres (Beton et al., 2022).

As for WT DNAJB1, we performed single-particle analysis using the amyloid fibre reconstruction toolbox in RELION 3.1 to study ΔJ-DNAJB1 in complex with αSyn amyloid fibres. The characteristic 4.8 Å amyloid repeat was visible in 2D classes and their corresponding FFT (Supplementary fig 6A,B). We then generated a structure at 3.2 Å (Supplementary Fig 6D and Supplementary fig 7) with a helical twist of −1.33° and a helical rise of 4.74 Å (Supplementary fig 6C). This structure is different from the two forms shown in Supp Figure 1, but similar to a reported structure with an additional density at the N-terminus displaying a hairpin-like shape (Supplementary fig 7, PDB: 6OSJ, Ni et al., 2019). However, we could not assign the sequence for the additional hairpin because of a gap in the density.

We then studied the complex with the ΔJ-DNAJB1 mutant. Like WT DNAJB1, micrographs of ΔJ-DNAJB1 bound to αSyn amyloid fibres showed a pseudo-regular binding of the chaperone (Fig 3A). 2D class averages and their power spectra showed the same 40 Å periodicity (Fig 3B). However, the pattern of decoration seen in projection was different from that of WT DNAJB1: instead of 2 layers of dots, ΔJ-DNAJB1 decoration was characterised by stripes perpendicular to the fibril axis (Fig 3B,C). Since the globular J-domain and G/F linker represent 46% of DNAJB1 molecular weight (17 kDa of the 37 kDa monomer), the electron density of the J-domain appears in projection as a bright dot in the 2D classes of the WT DNAJB1. We produced a structure of the complex at ∼22 Å, masked to include only one protofilament decorated with ΔJ-DNAJB1 (Fig 3D and Supplementary fig 8A,B). Like the complex with WT DNAJB1, ΔJ-DNAJB1 is positioned asymmetrically relative to the fibril core with one subunit closer to the fibre. The density corresponding to ΔJ-DNAJB1 was fitted with the crystal structure of DNAJB1 lacking the J-domain and the G/F linker, with a similar change in dimer angle to that in full length DNAJB1 (Fig 3D, PDB: 3AGY). As for WT DNAJB1 complex, we could not determine a structure at a higher resolution because the fibril core dominated the alignment. Lastly, we checked that reprojections of the map show a reasonable resemblance to the 2D classes, supporting the validity of the low-resolution map (Supplementary fig 8B,C).

**Figure 3.**
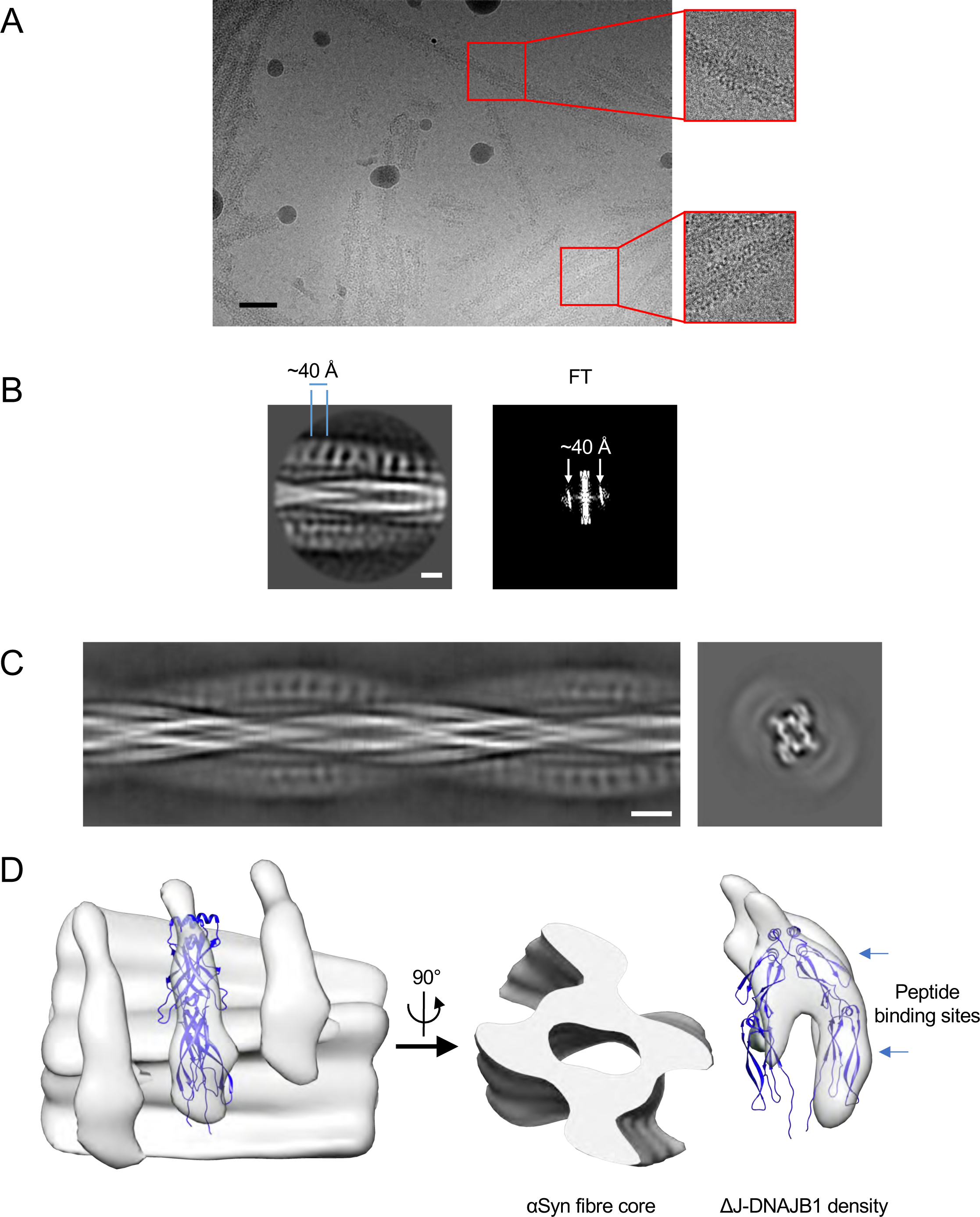
The complex of αSyn amyloid fibres and ΔJ-DNAJB1. A) Micrograph showing the fuzzy decoration of ΔJ-DNAJB1 on αSyn amyloid fibres. ΔJ-DNAJB1 repeats are discernible as dark dots. Scale bar, 50 nm. B) 2D class and FT showing the 40 Å repeat in ΔJ-DNAJB1 decoration, shown in reversed contrast. Scale bar, 5 nm. C) Side view of the aligned 2D classes and calculated cross-section. Scale bar, 100 Å. D) 3D reconstruction of the ΔJ-DNAJB1: αSyn fibre complex. The density corresponding to ΔJ-DNAJB1 was fitted with the crystal structure of the truncated human DNAJB1 lacking the J-domain and G/F linker (PDB: 3agy) using Flex-EM. Scale bar, 50 Å.

These structures were obtained using a buffer without salts (HD buffer), to enhance the amount of bound DNAJB1 to the fibres. Although the HD buffer was not physiological, it did not denature DNAJB1 and the stronger binding was reversed when the salts were added back. Earlier negative stain tomography of the complex at physiological salt concentration suggested a ∼50 Å repeat (Gao et al., 2015). Therefore, we conclude that the HD buffer did not affect the regularity of the binding but was useful to facilitate single particle analysis.

We attribute the limited resolution of the DNAJB1 maps, at ∼14 Å and ∼22 Å for WT DNAJB1 and ΔJ-DNAJB1 respectively, to the high flexibility of the 74 kDa DNAJB1 dimer in the complex (around 30 disordered residues separate the rigid core of the fibre from DNAJB1 density) that caused the alignment to be dominated by the highly ordered fibres. In the case of the ΔJ-DNAJB1 complex, the dimer is only 40 kDa, and the map had to be low pass filtered to retain the weak DNAJB1 electron density in the 2D class averages.

Despite the low resolution, the similarity in features between the WT DNAJB1 and ΔJ-DNAJB1 complexes, obtained completely independently with different fibre preparations, different DNAJB1 proteins and by different authors, suggests that the overall features are reliable. The ΔJ-DNAJB1 map shows that removal of the J-domain and G/F linker does not change the regular, asymmetric binding of DNAJB1 to the fibres.

### Helix V of the DNAJB1 J-domain is required to produce the organised, active clusters of chaperones at fibre ends

Since DNAJB1 J-domain and autoinhibitory sites are not required for the regular DNAJB1 binding to αSyn amyloid fibrils, we wondered whether the 2-step action of DNAJB1 could influence the pattern of Hsc70 recruitment. It has been shown that only class B JDPs can trigger fibre disaggregation, attributed to the additional auto-inhibitory helix V in the G/F region docking to the J-domain in a 2-step mechanism for Hsc70 recruitment to the substrate (Faust et al., 2020; Gao et al., 2015). Class A JDPs, which do not possess that additional helix, can recruit Hsc70 in a 1-step mechanism but are inactive in disaggregation. We used a mutant of DNAJB1 (ΔH5 DNAJB1), which contains 5 mutations (H99G, M101S, F102G, F105S and F106G) to generate an unstructured G/F region as in DNAJA2, and compared Hsc70 decoration between WT and ΔH5 DNAJB1. All the following experiments were performed in disaggregation buffer containing salts (Table 1) since both Hsc70 and Apg2 can precipitate in HD buffer (data not shown).

**Table 1:** Scoring of fibre decoration. Numbers in brackets and totals are numbers of fibre segments counted. Organised and disorganised categories both show dense binding.

|  | Organised dense | Disorganised dense | Total dense | Sparse | Not decorated | Not determined | Total |
| --- | --- | --- | --- | --- | --- | --- | --- |
| WT | 30% (42) | 2% (3) | 32% (45) | 40% (55) | 23% (31) | 5% (7) | 138 |
| ΔH5 | 7% (23) | 22% (67) | 29% (90) | 35% (108) | 32% (99) | 4% (13) | 310 |
| Total | 65 | 70 | 135 | 163 | 130 | 20 | 448 |

We first checked previously reported results by confirming with a ThT assay that ΔH5 DNAJB1 does not support the disaggregation of αSyn amyloid fibrils, unlike WT DNAJB1 (Fig 4A). Nonetheless, Hsc70 is recruited to similar levels (∼25% and 27%, respectively) in the presence of WT or ΔH5 DNAJB1 and Apg2, as verified by a binding assay (Fig 4B-D). We noted that Apg2 enhances Hsc70 binding with ΔH5 DNAJB1, as reported for WT DNAJB1 (Fig 4B,C, Beton et al., 2022).

**Figure 4.**
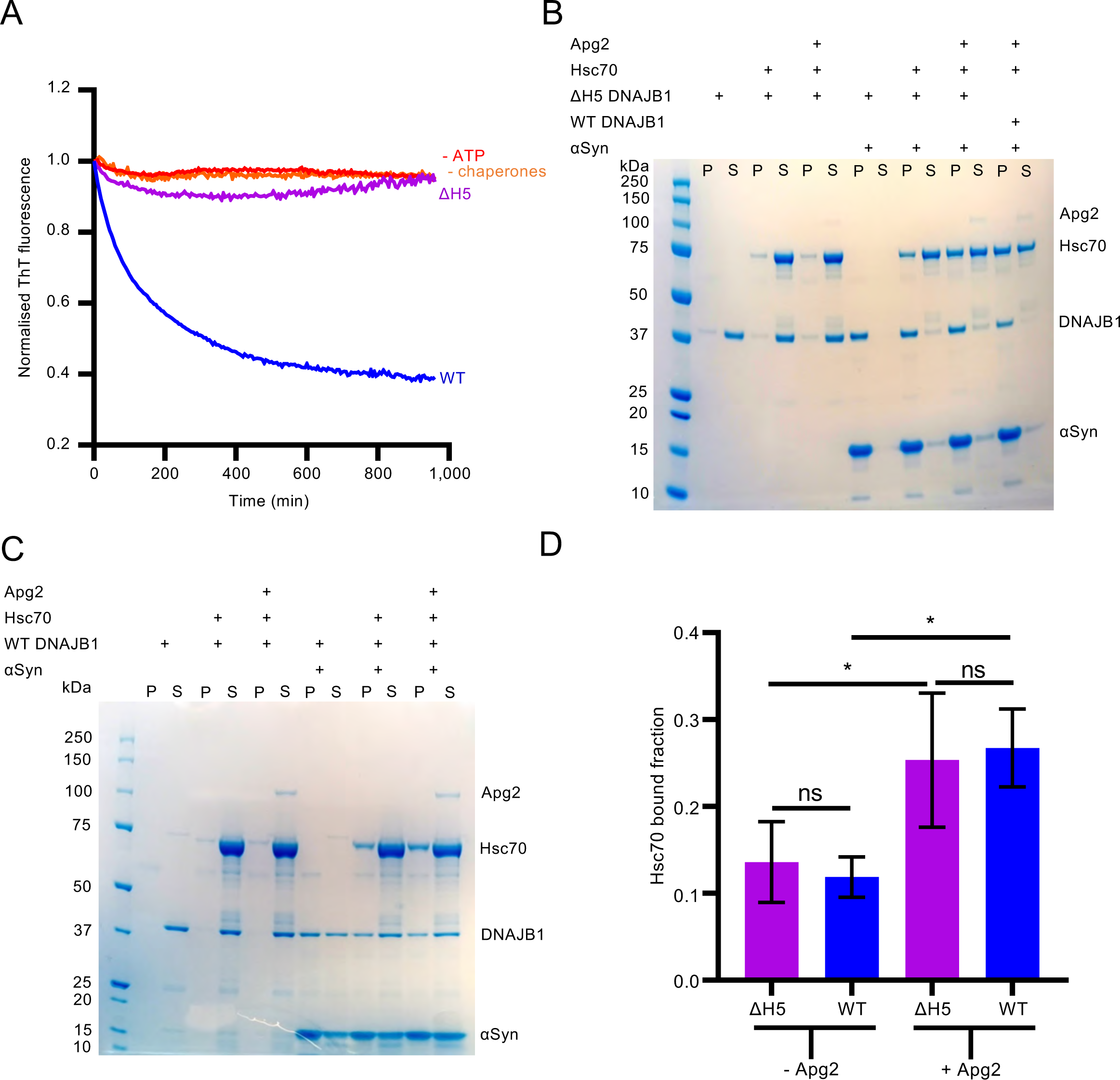
ΔH5 DNAJB1 recruits Hsc70 to the fibres at the same level as WT but is inactive in disaggregation. A) ThT assay showing that ΔH5 DNAJB1 cannot trigger the disaggregation of αSyn amyloid unlike WT DNAJB1. B) Binding assay showing that ΔH5 DNAJB1 can recruit Hsc70 at the same level as WT DNAJB1 in the presence of Apg2. ΔH5 DNAJB1 also enhances Hsc70 recruitment in the presence of Apg2. C) Binding assay showing that Apg2 can enhance Hsc70 recruitment in the presence of WT DNAJB1. D) Histogram of the Hsc70 recruitment to the amyloid fibrils as a function of Apg2 and WT or ΔH5 DNAJB1 in each condition shown in (B) and (C) (N = 3 independent experiments). A Shapiro-Wilk test was performed to check the normality of the data, followed by a one-way ANOVA with Tukey’s multiple comparisons test (* P= 0.0142, ** P = 0.0014). Mean Hsc70 binding values with SD are shown.

The normal level of Hsc70 recruitment combined with a complete loss of disaggregation activity led us to search for structural differences between the WT and mutant assemblies by cryo-electron tomography (cryo-ET). Complexes composed of fibrils, WT DNAJB1, Hsc70 and Apg2 show highly organised clusters (Fig 5A,B), whereas the complexes containing ΔH5 DNAJB1 have equally dense, but less regularly distributed decoration on the fibres (Fig 5C). The organised clusters contain densely packed globular features tethered to the fibres, extending radially outwards from the fibre surfaces along spiral tracks following the helical twist of the protofilaments with a spacing of ∼50-70 Å. The globular densities extend out to ∼140 Å from the fibre surface along the fibres, estimated by measuring the full outer diameter and assuming a fibre diameter of 100 Å, and often appear lined up in rows. For comparison, DNAJB1 density in the 2D class averages (Fig 2C; Supp. Fig 2) extends ∼80-100 Å from the fibre. In the mutant complexes, the recruited densities are scattered over all radii on the fibres. We quantified the differences in chaperone distribution between WT and ΔH5 complexes by blinded counting (Table 1). The ordered features were seen in 93% (42/45 of the densely bound segments) of the amyloid fibrils densely decorated with WT chaperones, whereas only 25% (23/90 dense segments) of the densely decorated fibrils display organised complexes in the presence of ΔH5 DNAJB1 (Table 1).

**Figure 5.**
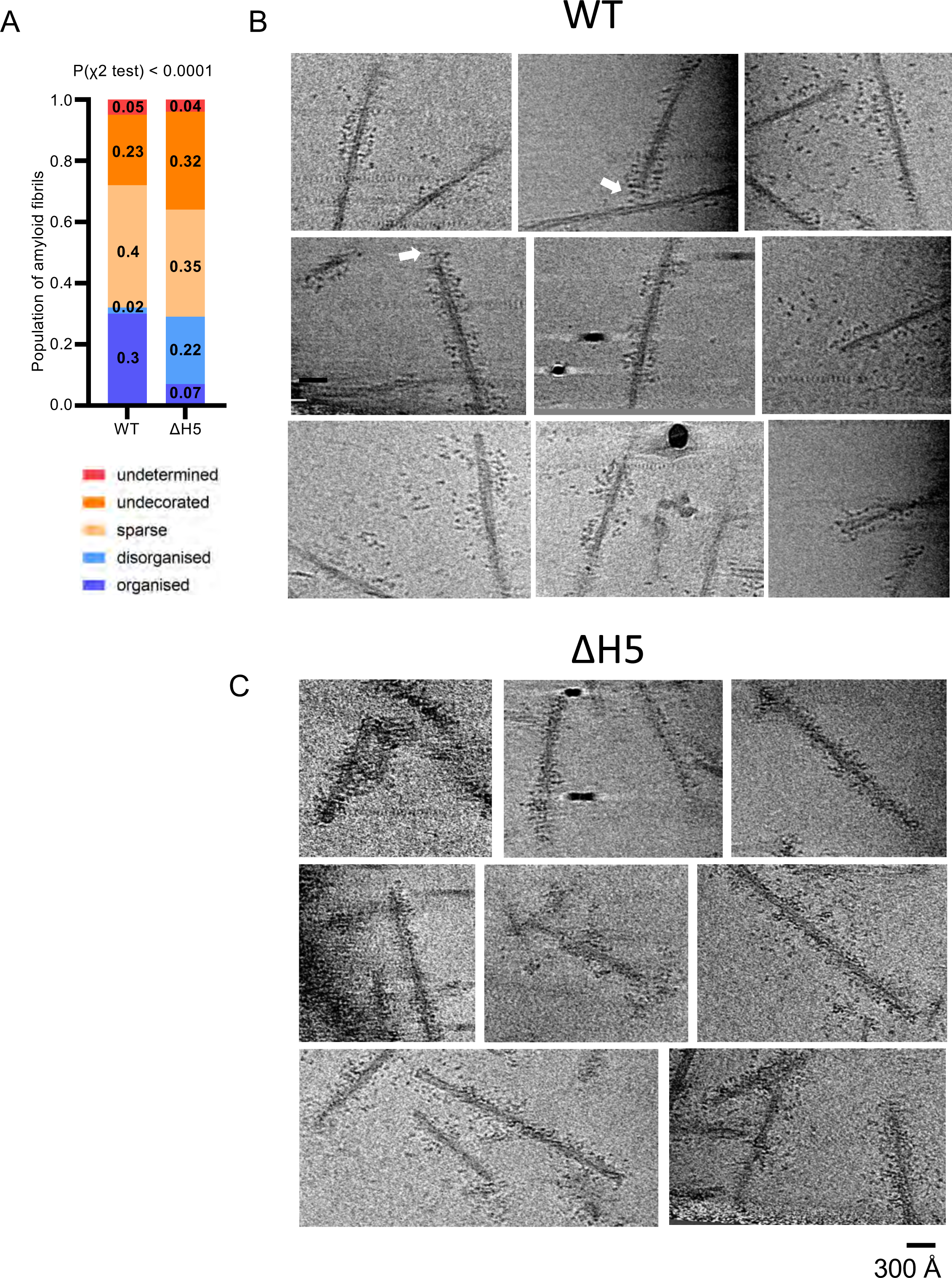
Structural comparison by cryo tomography of WT and ΔH5 DNAJB1 chaperone complexes on αSyn amyloid fibres. A) Plot of blinded counting of densely bound regions on the fibre segments, categorised as either organised or disorganised; sparsely bound regions, unbound, or undetermined. The fraction of organised regions is significantly higher in the WT complex (P(χ^2^ test) < 0.0001). Fractional values are shown in the plot and the data are tabulated in Table 1. B) Tomogram slices of examples of the WT complexes, showing the arrays of chaperones spiralling around the fibres, with rows of globular densities at the ends of stalks extending from the fibre surface. White arrows indicate examples of the chaperone decoration bending around a fibre end. The maximum diameter of the decorated fibres is ∼390 Å. C) Tomogram slices of ΔH5 DNAJB1complexes, showing a less organised binding, with the chaperones more irregularly distributed at different radii. The maximum diameter also reaches 390 Å, but much less consistently than in the WT system. All tomogram slices are averages of 5 sections. Scale bar, 300 Å.

## Discussion

In this study, we have probed the structural basis of amyloid fibre disaggregation by the Hsc70 chaperone system, beginning with single particle cryo-EM to visualise substrate engagement by the DNAJB1 co-chaperone with αSyn amyloid fibres. This revealed DNAJB1 binding in an unanticipated, asymmetric position with one subunit of the homodimer closer to the fibre surface, packed with a ∼40 Å repeat all along the fibres. This pattern of binding was observed for both WT and ΔJ-DNAJB1 in independent experiments with different fibre conformers. The binding is consistent with previous studies showing that i) DNAJB1 binds to the region of residues 123-129 of the disordered C-terminus of αSyn and ii) ΔJ-DNAJB1 can also bind to αSyn fibrils (Beton et al., 2022; Wentink et al., 2020). The approximate fit of DNAJB1 and αSyn (Fig 2 and Supplementary fig 4C) shows that the αSyn C-terminus exits the fibre core at a feasible position for αSyn residues 123 – 129 to contact the DNAJB1 substrate binding domain (Fig 2D; Wentink et al., 2020).

We next considered the role of the two-step auto inhibition of DNAJB1. Since the ΔH5 DNAJB1, mutated to remove autoinhibition by helix 5 in the G/F linker, supported full recruitment of Hsc70 to the fibres but not disaggregation activity, it was important to compare the structures of the fully assembled, active and inactive complexes on fibres by cryo ET. Previous AFM and TIRF studies showed short bursts of disassembly after a period of chaperone recruitment, usually at fibre ends, over a limited length of fibril, 1000-2000 Å (Beton et al., 2022). The tomograms show recruited Hsc70 as globular densities at some distance from the fibre surface. In the wild type system they appear to be held on stalks of density extending away from the fibre, following the helical symmetry of the fibres (Fig 5B). On the other hand, with the ΔH5 mutant, the recruited Hsc70 densities are scattered more irregularly over all radii on the fibres (Fig 5C). We propose that the organised trains of wild type chaperones constitute the hot spots being primed for the bursts of disaggregation previously observed by AFM (Beton et al., 2022).

In the attempt to build a structural model of the complex, we found that the fibre:DNAJB1 structure places strong steric constraints on Hsc70 binding. The tight packing of the J domains against the CTDs in theDNAJB1:αSyn fibre complex (Fig 2D) is consistent with the auto inhibited form, and docking of Hsc70 to the accessible J domain via the known Hsp70:J domain interface (eg PDB 5nro) results in severe clashes between the ATPase domain of Hsc70 and the CTDs of DNAJB1 (Supplementary Fig 9). However, activation of DNAJB1 by binding of the EEVD tail of Hsc70 to the CTD sites releases the J domain (Faust et al., 2020), so that it is tethered by a 40 residue flexible linker, allowing it to diffuse up to 120 Å away from the CTD. The Hsc70 can readily bind the released J domain on the outside of the DNAJB1 layer. This constraint allowed us to create a schematic model with the DNAJB1 dimer based on the structure in Fig 2D, with the outer J domain detached and an alpha fold model of Hsc70:DNAJB1 J domain placed arbitrarily on the outside of the complex, avoiding clashes (Fig 6A). To assess which features correspond to the DNAJB1 and Hsc70, the density projection of this model (Fig 6B) is compared to a tomogram section of the full WT complex (Fig 6C) and to the class average projection of DNAJB1:αSyn fibre (Fig 6D). Although the precise positions and orientations of the Hsc70 proteins are not known, the tomogram densities and steric constraints of the DNAJB1 clearly locate the bulk of the visible Hsc70 densities over 100 Å from the fibre surface. This leads to the unexpected conclusion that the rows of recruited Hsc70 proteins are too far away to reach their substrate binding sites on the N-termini of αSyn, which are either within 14 residues of the fibre surface (ie a maximum extent of ∼40 Å) or on the surface (Wentink et al., 2020). This leaves the exposed fibre end as the only site at which the Hsc70 can bind its substrate. The tomograms sometimes show the chaperone densities bending around at the fibre ends (white arrows, Fig 5B). In support of this finding, the previous AFM videos showed zones of dense chaperone binding on the fibres, which then unravelled from one end after a delay (Beton et al., 2022). Since the fibres are densely covered with DNAJB1 binding sites, the autoinhibition likely coordinates the assembly of the ordered trains of chaperones. These ordered trains could then produce a cascade of disassembly from the exposed fibre end.

**Figure 6.**
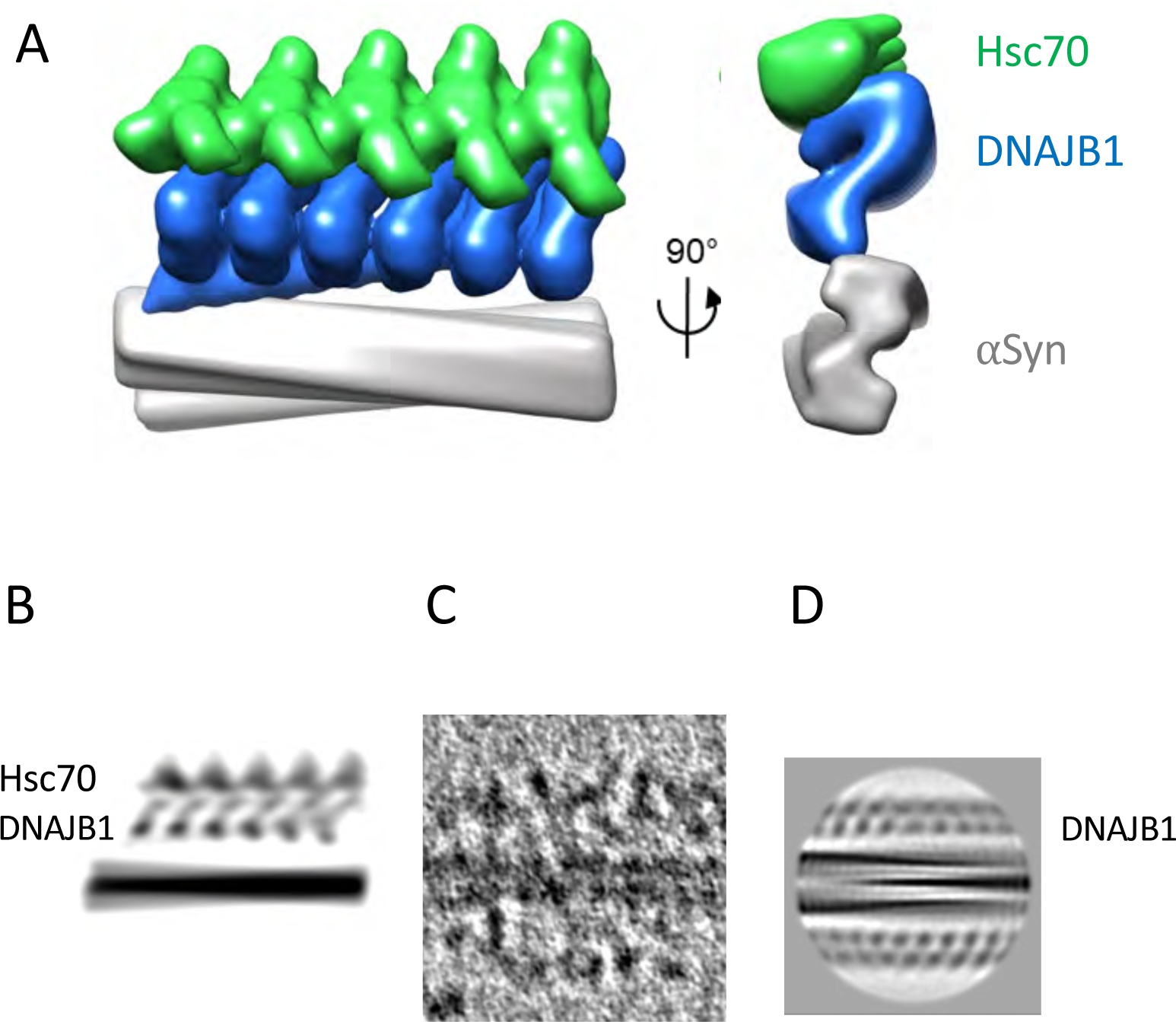
Model of Hsc70 binding and activation. A) A schematic model of Hsc70 binding to the αSyn:DNAJB1 complex. An alpha fold model of Hsc70:DNAJB1 J-domain was placed on the outside of the complex with the outer J-domain removed from the fitted DNAJB1 model, avoiding clashes. The Hsc70 orientation is arbitrary but it is not possible to place it any closer to the αSyn N termini, because of the presence of the DNAJB1. The model is shown in side and end views. Hsc70 is too bulky to pack every 40 Å as for DNAJB1 and is spaced at 50 Å. The 40 residue long flexible linker to the activated J domain can accommodate a wide range of positions for the Hsc70. B) Projected density of the model for comparison to the tomogram section in C). C) Tomogram section of the full chaperone system bound to an αSyn fibre. D) Class average of αSyn:DNAJB1 reversed in contrast from Fig 2 (protein density dark) for comparison. The approximate locations of DNAJB1 and Hsc70 layers are indicated with text labels. Panels B-D are shown approximately to the same scale.

The regions of ordered binding suggest a model, presented as a schematic cartoon in Fig 7, for the 2-step mechanism in which coordinated binding and activation of Hsc70s are favoured by intermolecular interactions between DNAJB1 dimers. The visible blobs on stalks arrangement is consistent with binding of the Hsc70 EEVD motif to the C-terminal domain I (CTD-I) (Wentink et al., 2020) of the more accessible, outer DNAJB1 subunit, releasing the auto-inhibitory helix V from the J-domain. This releases the J-domain to activate an Hsc70 by binding its ATPase domain (Faust et al., 2020). To form the ordered regions, we propose that the freed J-domain activates a second Hsc70 bound to an adjacent DNAJB1, which in turn could activate its neighbour, as suggested in Figure 7. The J-domain binding site of DNAJB1 and the C-terminal EEVD site of Hsc70 (which is at the end of a 30-residue disordered tail) can easily bridge between the corresponding binding sites on adjacent DNAJB1 dimers. In contrast, the ΔH5 mutant, acting like a class A JDP, allows uncoordinated binding to all sites and does not produce any disassembly.

**Figure 7.**
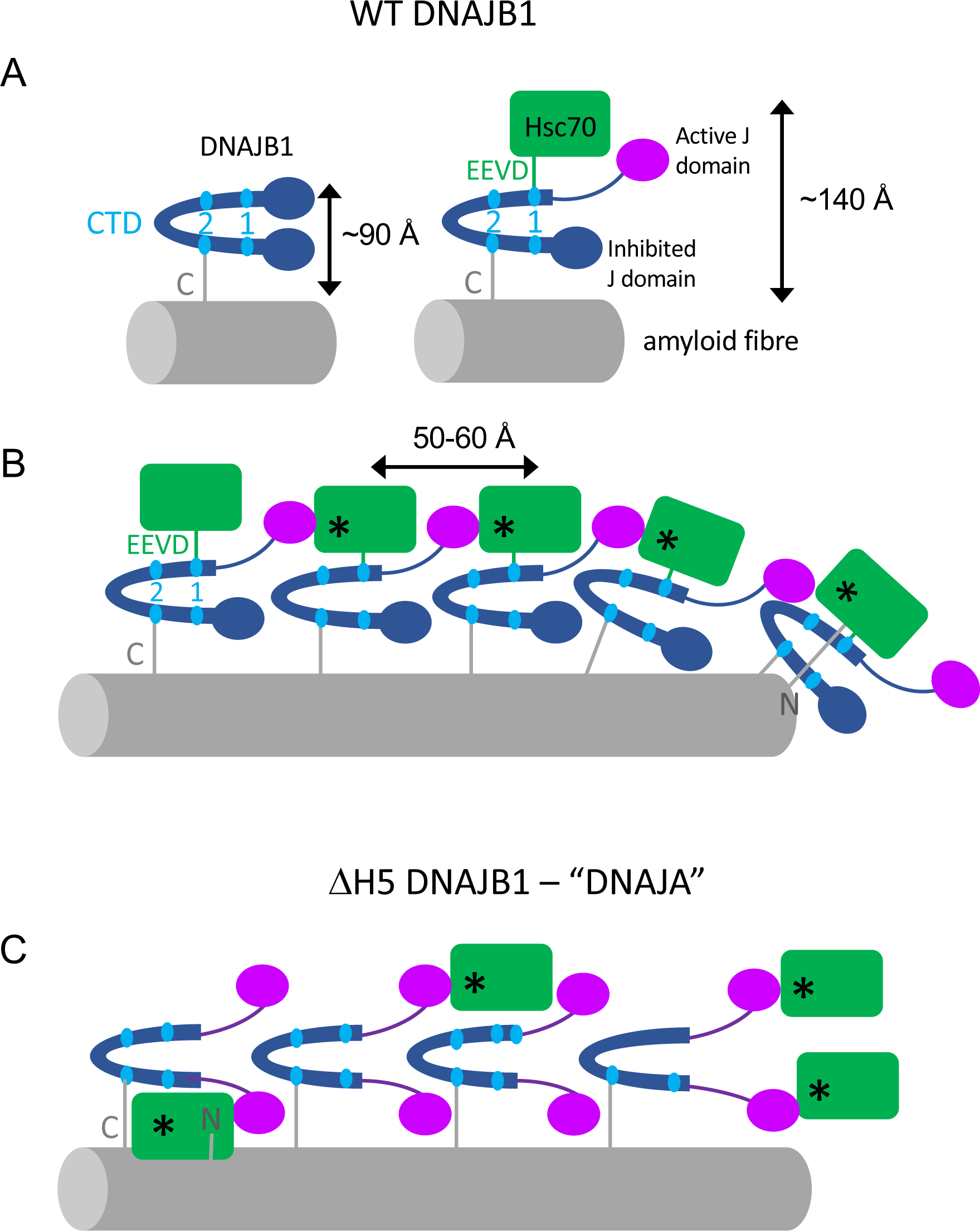
Cartoon showing proposed cooperative, 2-step mechanism of JDP recruitment of Hsc70 into active clusters for disaggregation. A) WT DNAJB1 binds to the flexible C-terminus of αSyn via its CTD2 site. Then, the EEVD motif of Hsc70 binds CTD1 and releases the J domain, which is proposed to activate an Hsc70 (*) on the adjacent DNAJB1 dimer. This could create a localised cloud of consecutive, actively recycling Hsc70s corresponding to the ordered arrays seen by cryo ET. B) The Hsc70s are packed with a spacing of ∼50-60 Å along the fibre and extend out >∼100 Å from the fibre surface, suggesting that they bind preferentially to the outer subunit of DNAJB1. Only the Hsc70 in complex with DNAJB1 at the fibril end can approach closely enough to interact with the N-terminus of αSyn. The binding follows the helical path of the fibre structure, omitted here for clarity. ATPase cycling is catalysed by Apg2, which is also omitted from the cartoon. It is present at up to 10% of the amount of Hsc70 but is not seen bound to the fibres in the binding assays. C) In the ΔH5 mutant system mimicking a class A JDP, all J-domains are already released. Therefore, Hsc70 appears to bind more randomly at all radii, suggesting that it can bind with equal probability to either DNAJB1 subunit. The resulting complexes are not active in disaggregation.

The 2-step mechanism could trigger a cascade of Hsc70 recruitment and activation, leading to the coordinated formation of organised, dense clusters of chaperones, which, along with ATPase cycling stimulated by transient binding of Apg2, are required for the disaggregation of amyloid fibrils (Beton et al., 2022; Wentink et al., 2020).

In summary, we have presented the first structures of a chaperone bound to amyloid fibrils and the first views of the differing effects of class A and B JDPs at a molecular scale. It remains a future challenge to verify the cooperative formation of organised clusters by class B JDPs and Apg2 and the mechanism of the unidirectional bursts of disaggregation.

## Acknowledgements

We thank Natalya Lukoyanova and Shu Chen for EM support, David Houldershaw for computing support and Claire Bagneris for molecular biology support. We are grateful to Rina Rosenzweig for the gift of mutant JDPs, and to Euan Pyle, Guendalina Marini and Jesus Gomez for advice and discussions. For comments on the manuscript we thank Bart Hoogenboom, Bernd Bukau and Rina Rosenzweig.

Most of the cryo-EM data were collected at the ISMB EM facility at Birkbeck College, University of London with financial support from the Wellcome Trust (202679/Z/16/Z and 206166/Z/17/Z). We acknowledge Diamond Light Source for access and support of the cryo-EM facilities at the UK’s national Electron Bio-imaging Centre (eBIC) [under proposal EM 20287], funded by the Wellcome Trust, MRC and BBRSC. The work was supported by a Wellcome Trust Investigator award to HRS (106249/Z/14/Z), a Wellcome Trust studentship to JB, and a Birkbeck Anniversary studentship to JM.

## Data availability

The cryo EM reconstructions and corresponding atomic models are deposited in the EMDB and PDB with the accession codes EMD-19443, 19444, 19448, 19441, 19462, 19461, 19446, 19447 and PDB 8RRR, 8RQM.

Raw data for binding and disaggregation assays are at https://researchdata.bbk.ac.uk/id/eprint/326/ and https://researchdata.bbk.ac.uk/id/eprint/328/ respectively.

## Author contributions

JM, JGB and ECJ carried out the experimental work; all authors contributed to analysing the data and writing the paper. HRS, JGB and JM designed research.

## Materials and Methods

### Protein purification

WT proteins (αSyn, DNAJB1, Hsc70, Apg2) were purified as described previously (Beton et al., 2022). Briefly, αSyn was expressed in DE3 Star *E. coli* cell lines (Thermo Fisher Scientific). Cells were lysed and the soluble fraction was boiled for 20 minutes, treated with streptomycin sulphate and ammonium sulphate before being dialysed overnight into deionised water. The dialysed sample was then purified by ion exchange chromatography with a HiTrap Q HP column followed by further size exclusion chromatography (SEC). The WT chaperones (DNAJB1, Hsc70, Apg2) were expressed in *E. coli* Rosetta^TM^ DE3 cells with a His_6_-Smt_3_ tag at the N-terminus and purified by nickel affinity chromatography. An additional SEC step was performed for DNAJB1.

The purified mutants of DNAJB1 (ΔJ-DNAJB1, the mutant lacking the J-domain and the G/F linker and ΔH5 DNAJB1, mutation of the helix 5 inhibitory binding site: H99G, M101S, F102G, F105S and F106G) were kindly provided by Rina Rosenzweig’s group (Weizmann Institute). Briefly, the mutants were purified by ion metal affinity chromatography (IMAC, Nickel), dialysed and cleaved with TEV protease, subjected to reverse IMAC and then subjected to SEC. The proteins were stored in 25 mM HEPES pH 7.5, 200 mM KCl, 10 mM MgCl_2_, 2 mM DTT.

### αSyn fibrillation reaction

αSyn amyloid fibres were produced as previously described (Beton et al., 2022; Gao et al., 2015; Nachman et al., 2020): monomeric αSyn (200 µM, 1 mL total volume) was incubated in fibrillation buffer (50 mM NaPO_4_, 100 mM NaCl, 0.05% w/v NaN_3_, pH 7.30) in a 1.5 mL Protein LoBind Eppendorf tube (Eppendorf, Germany) on an orbital shaker (PCMT Thermoshaker Grant-bio) at 1,000 rpm, 37°C for 1 week. The formation of amyloid fibres, rather than amorphous aggregates, was confirmed by negative stain electron microscopy.

### Binding assay

αSyn amyloid fibres were sonicated for 15 min at high frequency using a CPX 2800 Bransonic Ultrasonic bath (Branson) to maintain them dispersed and chaperone aliquots (WT, ΔJ and ΔH5 DNAJB1) were centrifuged at 17,000g for 30 minutes at 4°C to discard any chaperone aggregates. αSyn fibres (20 µM, monomer concentration) were incubated with DNAJB1 (8 µM) in HD or HKMD buffer (table 1) for 30 minutes at 30°C. The samples were then centrifuged at 17,000g for 30 minutes. The supernatant was collected and the two fractions (pellet and supernatant) were incubated in 4X NuPAGE LDS Sample buffer (Thermo Fisher Scientific) for at least 30 min at 90°C. Samples were then loaded on BOLT 4– 12% Bis-Tris gels (Thermo Fisher Scientific) and proteins were separated by SDS-PAGE. At least three independent experiments were performed for each condition. The statistical analyses were performed in Prism 8 (GraphPad).

Regarding the reversibility of the binding, a similar protocol was performed with 2 consecutive 30-minute incubations at 30°C in the buffer indicated in the figure. At least three independent experiments were performed for each condition. The statistical analyses were performed in Prism 8 (GraphPad).

### Thioflavin T (ThT) assay

αSyn amyloid fibres were sonicated for 15 min at high frequency using a CPX 2800 Bransonic Ultrasonic bath (Branson). αSyn fibres (2 µM, monomer concentration) were incubated with DNAJB1 (2 µM, WT or ΔH5 DNAJB1 when indicated), Hsc70 (4 µM), Apg2 (0.4 µM) and ThT (15 µM) in the disaggregation buffer (table 2). ThT fluorescence was recorded every 5 minutes for 16 h on a FLUOstar Omega plate-reader (BMG LABTECH, excitation: 440 nm, emission: 482 nm). Background ThT fluorescence of chaperones and buffer was subtracted and ThT fluorescence was normalised to the fluorescence intensity of the first time point (t = 0 min). At least three independent experiments were performed for each condition.

**Table 2.** The buffers used in this study.

| Buffer | Composition |
| --- | --- |
| HKMD buffer | 50 mM HEPES, 50 mM KCl, 5 mM MgCl<sub>2</sub>, 2 mM DTT, pH 7.50 |
| HD buffer | 50 mM HEPES, 2 mM DTT, pH 7.50 |
| Disaggregation buffer | HKMD buffer, 5 mM ATP, 6 mM PEP, 20 ng/μL pyruvate kinase |

Regarding the ThT assay monitoring the effects of the HD buffer on DNAJB1 activity, αSyn amyloid fibres and DNAJB1 were initially incubated in HKMD or HD buffer for 30 minutes at 30°C before performing the ThT assay in the disaggregation buffer in the presence of Hsc70 and Apg2.

### Cryo-EM Sample preparation

The reconstructions of DNAJB1:αSyn presented here were generated from 3 separate datasets. The details for grid preparations in each sample are outlined in table 3. In all cases, prior to grid preparation, αSyn fibres (200 μM monomer concentration) were sonicated for 45 minutes using a Branson 1800 Cleaner or for 15 minutes at high frequency using a Branson CPX 2800 Ultrasonic bath.

**Table 3.** Details of cryo-EM grid preparation.

| **Datasets** | **WT DNAJB1 dataset 1** | **WT DNAJB1 dataset 2** | **WT DNAJB1 dataset 3** | **ΔJ-DNAJB1 dataset** |
| --- | --- | --- | --- | --- |
| **αSyn fibre concentration (μM)** | 10 | 10 | 6 | 6 |
| **DNAJB1 concentration (μM)** | 20 | 20 | 2.4 | 2.4 |
| **Grid type** | C-flat 2/2 | C-flat 2/2 | C-flat 2/2, 400 mesh | C-flat 2/2, 400 mesh |
| **Grid preparation machine** | Vitrobot | Vitrobot | Leica EM GP2 | Leica EM GP2 |

### Cryo-EM Data Collection

The microscopes and imaging parameters used for collection of each dataset for single particle analysis are summarised in table 4.

**Table 4.** Details of cryo-EM data collection.

| **Datasets** | **WT DNAJB1 dataset 1** | **WT DNAJB1 dataset 2** | **WT DNAJB1 dataset 3** | **ΔJ-DNAJB1 dataset** |
| --- | --- | --- | --- | --- |
| **Microscope** | FEI Titan Krios | FEI Titan Krios | FEI Titan Krios | FEI Titan Krios |
| **Acceleration voltage (kV)** | 300 | 300 | 300 | 300 |

**Table 4. Details of cryo-EM data collection.**
| Camera | Gatan K2 | Gatan K2 | Gatan K3 | Gatan K3 |
| --- | --- | --- | --- | --- |
| Energy filter slit width (eV) | 20 | 20 | 20 | 20 |
| Super-resolution pixel size (Å) | 0.5025 | 0.5025 | 0.55 | 0.53 |
| Total frames | 50 | 50 | 50 | 50 |
| Total dose (e-/Å<sup>2</sup>) | 44 | 44 | 48.5 | 49 |
| Defocus range | 1.0 – 3.5 | 1.0 – 3.5 | 1.5 – 3.5 | 1.5 – 3.0 |
| Number of movies collected | 3261 | 2797 | 13429 | 9,943 |

### Helical Image Processing

All super-resolution movies were gain corrected, binned by a factor of 2, motion corrected and dose weighted using RELION’s CPU implementation of MotionCor2 (Zheng et al., 2017; Zivanov et al., 2018). Aligned, non-dose weighted micrographs were used for CTF estimation using CTFFIND4.1 (Rohou and Grigorieff, 2015). Visual inspection of real-space images and their power spectra was used to identify and discard images showing ice contamination or other defects.

### αSyn fibres from WT DNAJB1 datasets

Fibres were manually picked from images from dataset 1 and 2 (tables 2 and 3) in RELION 3.1 using the helical picking tool and particles were extracted as 320-pixel (336 Å) boxes along the fibre axis with an inter-box distance of 36 pixels (38.4 Å). Reference free 2D classification was used to identify two polymorphs (polymorphs 1 and 2) based on the approximate thickness and helical twist of fibres in 2D classes. These classes were then used to generate an initial model using the relion_helix_inimodel2d tool, with a helical rise of 4.8 Å for both classes and crossover distances estimated from the raw micrographs. This initial model was lowpass filtered to 10 Å prior to 3D refinements. Reconstructions were generated by multiple rounds of 3D auto-refinement and CTF-refinement in RELION. The final reconstructions were sharpened using RELION postprocessing and the resolution was estimated by Fourier shell correlation (FSC) at 0.143 between the half-maps. The polymorph 2 map was fitted with PDB:6RT0 in Chimera as a rigid body and refined using PHENIX (Adams, 2018; Pettersen et al., 2004).

### αSyn:WT DNAJB1 complex

Particles were picked from images in dataset 3 (Tables 2 and 3) using the filament picking mode in crYOLO 1.5.6. The deep learning network was trained using coordinates manually assigned from 60 micrographs using EMAN2 (Tang et al., 2007), and this trained network was used to pick particles from the remaining micrographs. 2,300,000 particles were picked using a 420-pixel box size (462 Å) with an inter-box distance of 80 Å. These particles were subjected to multiple rounds of 2D classification and, again, classes corresponding to each αSyn fibre polymorph were separated and processed individually. Initial models were generated using relion_helix_inimodel2d, with the crossover distances used for the fibre only reconstructions. 3D classification was used to identify sets of particles with DNAJB1 density at the fibre edge. Discernible DNAJB1 density was only observed for any classes of the fibre 2 polymorph, and further image processing on the fibre 1 subset was not done. For fibre 1 particles, C2 symmetry expansion was used to increase the effective number of selected particles. These particles were used to generate reconstructions through two rounds of 3D refinement in RELION: firstly, a consensus refinement without applying a mask, followed by a second round of refinement using a mask that incorporated the entire αSyn fibre and the DNAJB1 decoration along only a single protofilament. This reconstruction was sharpened using RELION postprocessing using a user supplied B-factor of −150 Å^2^. A predicted model of the DNAJB1 dimer was generated from the full sequence (taken from UNIPROT entry P25685 using ColabFold (Mirdita et al., 2022). This model was manually placed in density using Chimera (Pettersen et al., 2004) and flexibly refined into the density using Flex-EM (Topf et al., 2008). RIBFIND restraints (Malhotra et al., 2023) were generated manually for each domain of DNAJB1 (CTD-I, CTD-II and the J-domain) to prevent overfitting of the model (Joseph et al., 2016; Jumper et al., 2021; Mirdita et al., 2022).

### αSyn fibres from the ΔJ-DNAJB1 dataset

Particles were picked from images in ΔJ-DNAJB1 dataset 3 (tables 2 and 3) using the filament picking mode in crYOLO 1.5.6 (Wagner et al., 2019). The deep learning network was trained from 100 micrographs that were manually picked in crYOLO. 911,478 particles were picked using a 420-pixel box size (445 Å) with an inter-box distance of 44.5 Å, binned by a factor 3.3 to yield 128-pixel boxes. These particles were subjected to three rounds of 2D classification. Only particles showing decorated fibres were selected and the corresponding particles were re-extracted without binning and subjected to two additional reference-free 2D classifications. The classes showing the 4.8 Å cross-β repeat were selected to produce an initial model using the relion_helix_inimodel2d tool, with a helical rise of 4.8 Å and a crossover distance estimated at 620 Å from the raw micrographs. Reconstructions were generated by multiple rounds of 3D auto-refinement and CTF-refinement in RELION. The final reconstruction was sharpened using a B-factor of −84 Å^²^ in RELION postprocessing. The resolution was estimated from FSCs at 0.143 between the half-maps. The map was fitted with PDB: 6OSJ in Chimera as a rigid body, manually refined in COOT and automatically refined in PHENIX (Emsley and Cowtan, 2004).

### αSyn:ΔJ-DNAJB1 complex

The 3.3x-binned particles were subject to 3 consecutive 2D classification without restricting translation along the fibril axis. Best 2D classes were used to generate an initial model using the relion_helix_inimodel2d tool, with a crossover distance estimated at 620 Å. A round of 3D classification was followed by two rounds of 3D auto-refinement. A mask to retain the entire αSyn fibre and only the ΔJ-DNAJB1 decoration along a single protofilament was generated and used for the second 3D refinement. An ad-hoc 25 Å low-pass filter and a B-factor of −196 Å^2^ were applied in the postprocessing stage. The predicted model of DNAJB1 (generated as described above) was truncated (J-domain and G/F linker residues were removed) and manually placed in the density using Chimera. The model was then flexibly refined using Flex-EM with RIBFIND restraints for each domain.

### Cryo-ET Sample Preparation

Prior to grid preparation αSyn amyloid fibres were sonicated on high frequency for 30 min using a Branson CPX 2800 ultrasonic bath to produce dispersed fragments of αSyn fibres. The suspension was diluted to 6 µM αSyn monomer concentration and mixed with 6 µM Hsc70, 3 µM DNAJB1 (ΔH5 or WT) and 0.6 µM Apg2 and incubated for 1 h at 30°C in disaggregation buffer. 4 μL of the wild type preparation were applied to negatively glow discharged holey carbon C-flat grids (CF-1.2/1.3-4C) (Protochips, USA). 4 μL of the ΔH5 preparation were applied to negatively glow discharged holey carbon C-flat grids (CF-2/2-3Cu-50) (Protochips, USA). Both sets of grids were then back blotted before adding 3 μL of 10 nm Protein-A-coated gold (EMS, USA) as fiducial markers for 3D reconstruction of tilt series. The grids were then back blotted a second time before plunge freezing in liquid ethane using a Leica EM GP2 (Leica Microsystems, Germany)

### Cryo-ET Data Collection and Tomogram Reconstruction

The microscopes and imaging parameters used for collection of each dataset for tilt series are summarised in table 6. Tilt series were acquired from +60° to −60° with a 3° increment in a dose symmetric acquisition scheme.

**Table 5.**
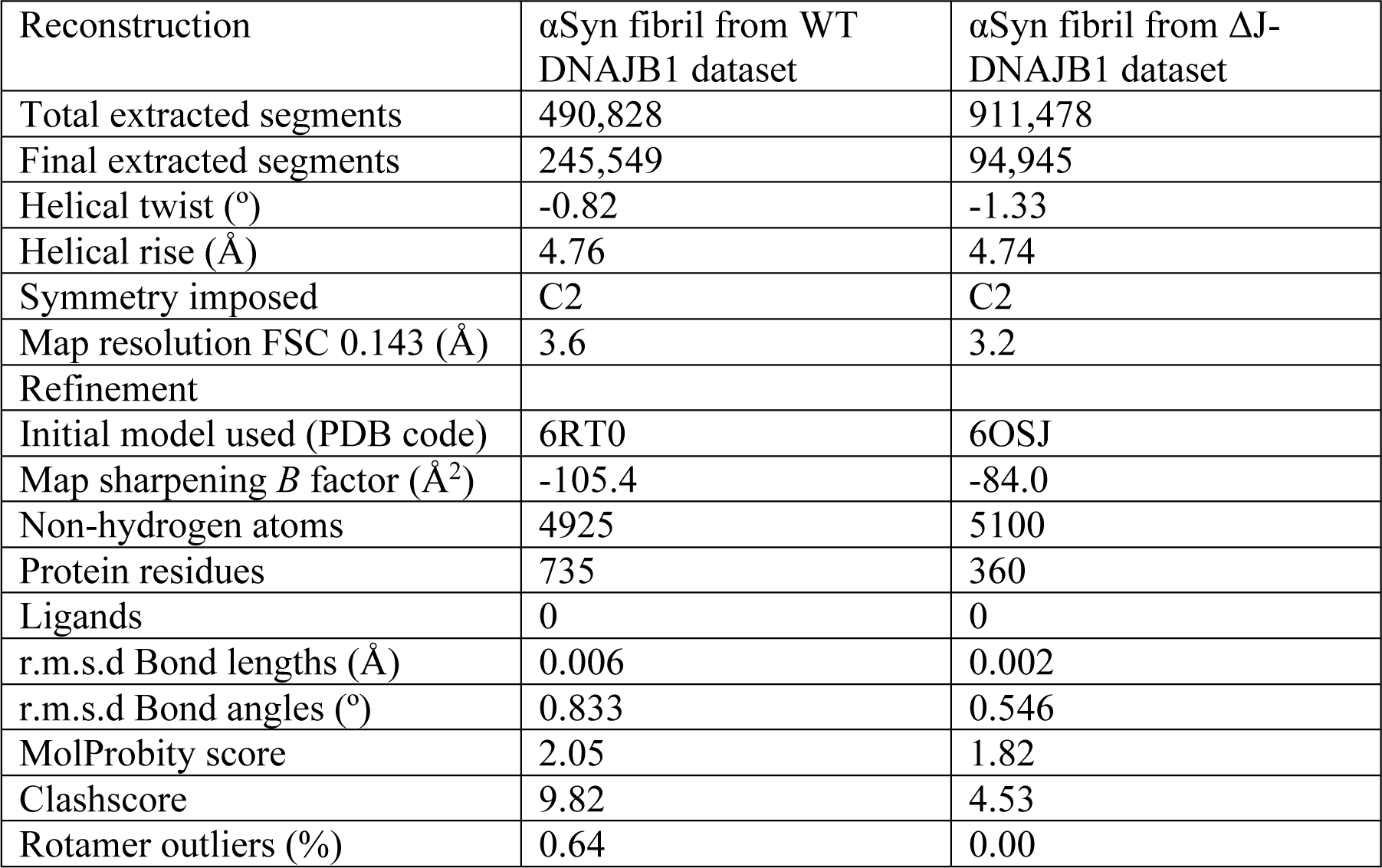
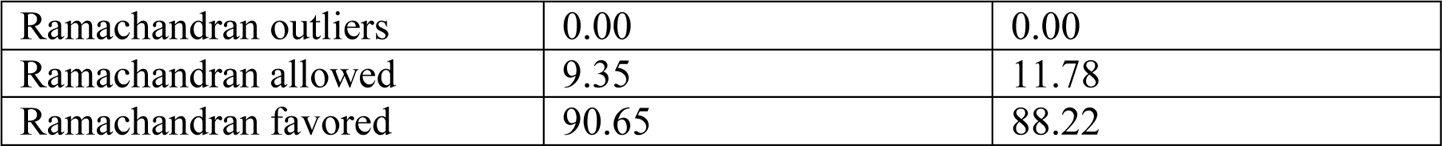
Details of cryo-EM data processing and model refinement.

**Table 6:** Details about tilt series data collection.

| Datasets | WT DNAJB1 dataset 1 | WT DNAJB1 dataset 2 | ΔH5 DNAJB1 |
| --- | --- | --- | --- |
| Microscope | FEI Titan Krios III | FEI Titan Krios III | FEI Titan Krios III |
| Acceleration voltage (kV) | 300 | 300 | 300 |
| Camera | Selectris Falcon IV | Selectris Falcon IVi | Selectris Falcon IVi |
| Energy filter slit width (eV) | 5 | 3 | 3 |
| Pixel size ($\text{\AA}$) | 1.9 | 1.94 (rounded to 1.9) | 1.94 (rounded to 1.9) |
| Total dose ($\text{e}/\text{\AA}^2$) | 118 | 116 | 116 |
| Defocus range ($\mu\text{m}$) | -2 to -6.5 | -2 to -4.4 | -2 to -4.4 |
| Number of movies collected | 3261 | 2797 | 13429 |

## Reconstruction method

For all tomography datasets, frames underwent whole frame alignment in MotionCor2 version 4.1 (Zheng et al., 2017) and defocus estimation was done using CTFfind version 4.1 (Rohou and Grigorieff, 2015). Tomograms were not dose weighted or CTF corrected. Tilt series alignment for WT DNAJB1 dataset 1 was performed using Dynamo version 1.1.333 (Castano-Diez et al., 2012). Tilt series alignment for WT DNAJB1 dataset 2 and for ΔH5 DNAJB1 were performed using fiducial based alignment in IMOD version 4.9.0 (Kremer et al., 1996). For all tilt series from WT DNAJB1 dataset 1, motion correction CTF estimation and tilt series alignment were automated using scripts and the guide from the following GitHub repository (https://github.com/EuanPyle/relion4_tomo_robot). For all tilt series alignments from WT DNAJB1 dataset 2 and ΔH5 DNAJB1, motion correction CTF estimation and tilt series alignment were automated using a custom build of the tomography module of Relion 4. All tomograms were binned by 2 in X and Y.

## Counting of fibres according to type of chaperone decoration

Chaperone binding along the fibres appeared either dense or sparse, and the densely bound regions appeared either organised, with the bound features spaced at roughly 60 Å, or disorganised, with no discernible regularity or repeat. In order to quantitate these observations, the tomograms of wild type and ΔH5 complexes were randomly mixed and provided without identification for blind counting by one author (Figure 5). This was done by searching through the planes of each tomogram to find isolated fibres, not in contact with other structures, that could be scored according to the nature of the decoration.

---

**Supplementary figure 1.**
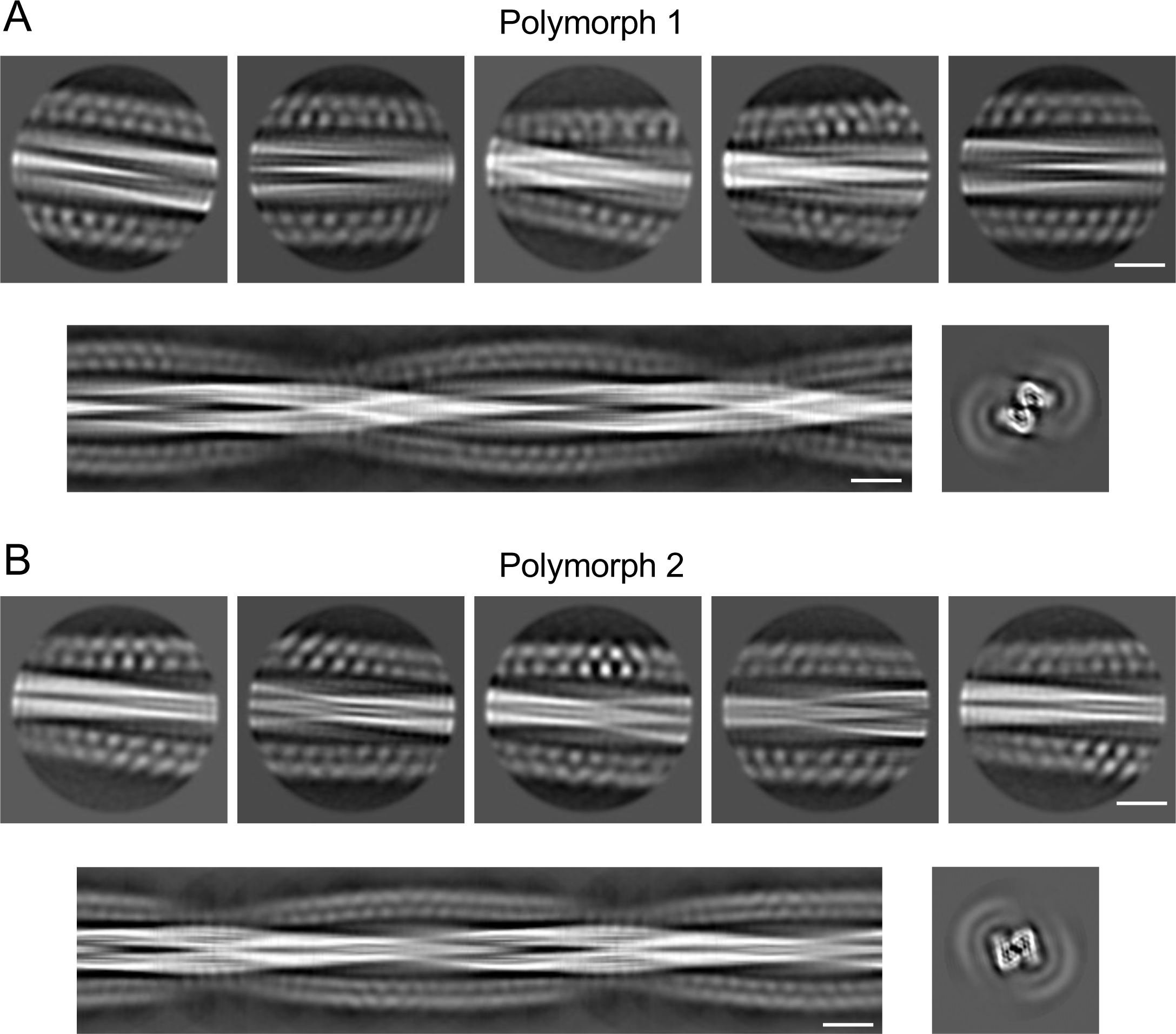
Diversity of fibre conformers. 2D classes, side view of the aligned 2D classes and calculated cross-section for the first (A) and second (B) polymorphs. Scale bars, 100 Å.

**Supplementary figure 2.**
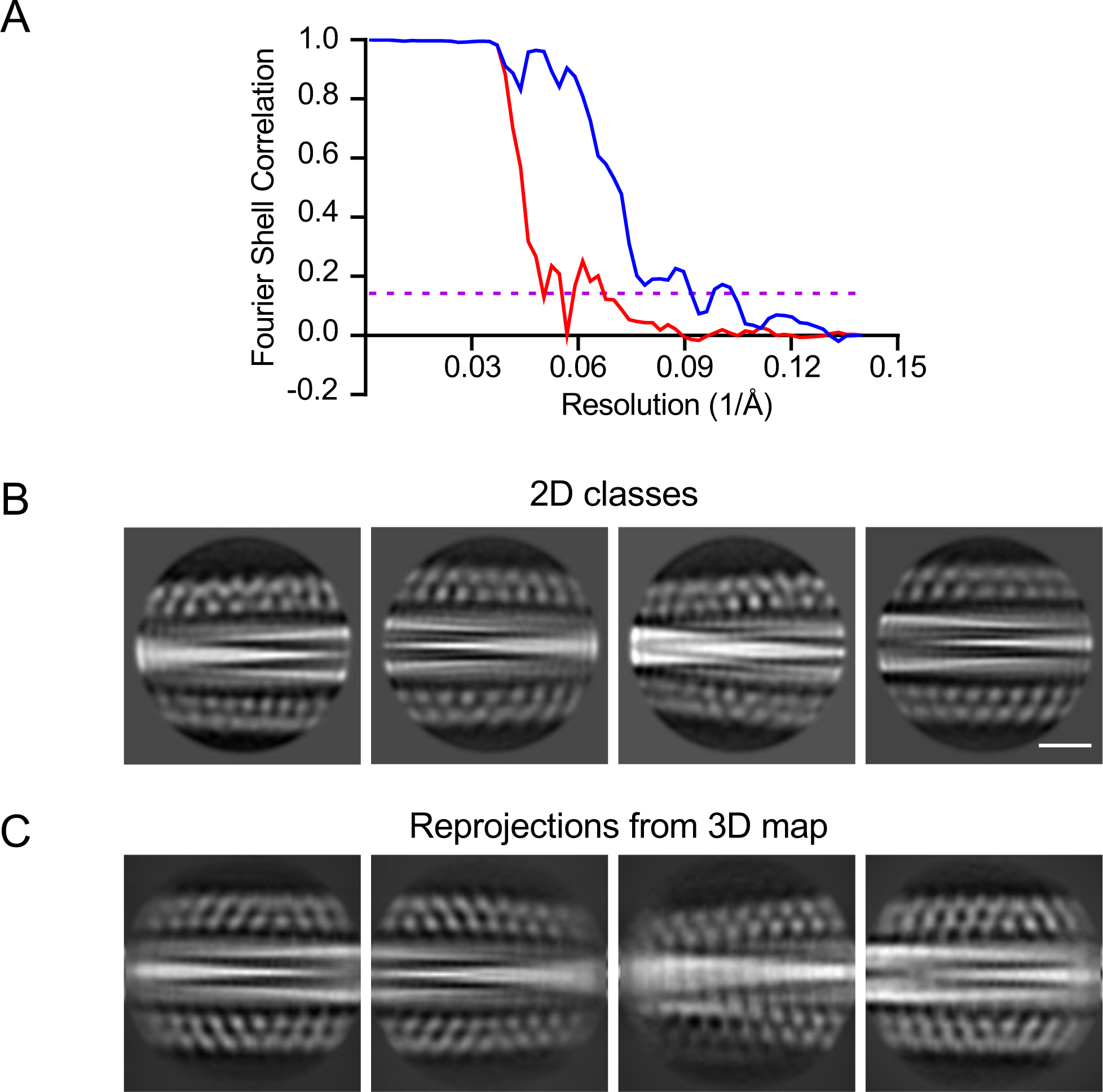
Resolution of WT DNAJB1:αSyn fibre complex map, 2D classes and reprojections from the map. A) FSC curves of two independently refined half-maps showing phase randomisation (red curve) and the cryo-EM reconstruction (blue curve). The dashed purple line indicates FSC = 0.143 B) Additional 2D classes showing the decorated αSyn fibres. Scale bar, 100 Å. C) Reprojections calculated from the 3D map. The reprojections resemble the 2D class averages.

**Supplementary figure 3.**
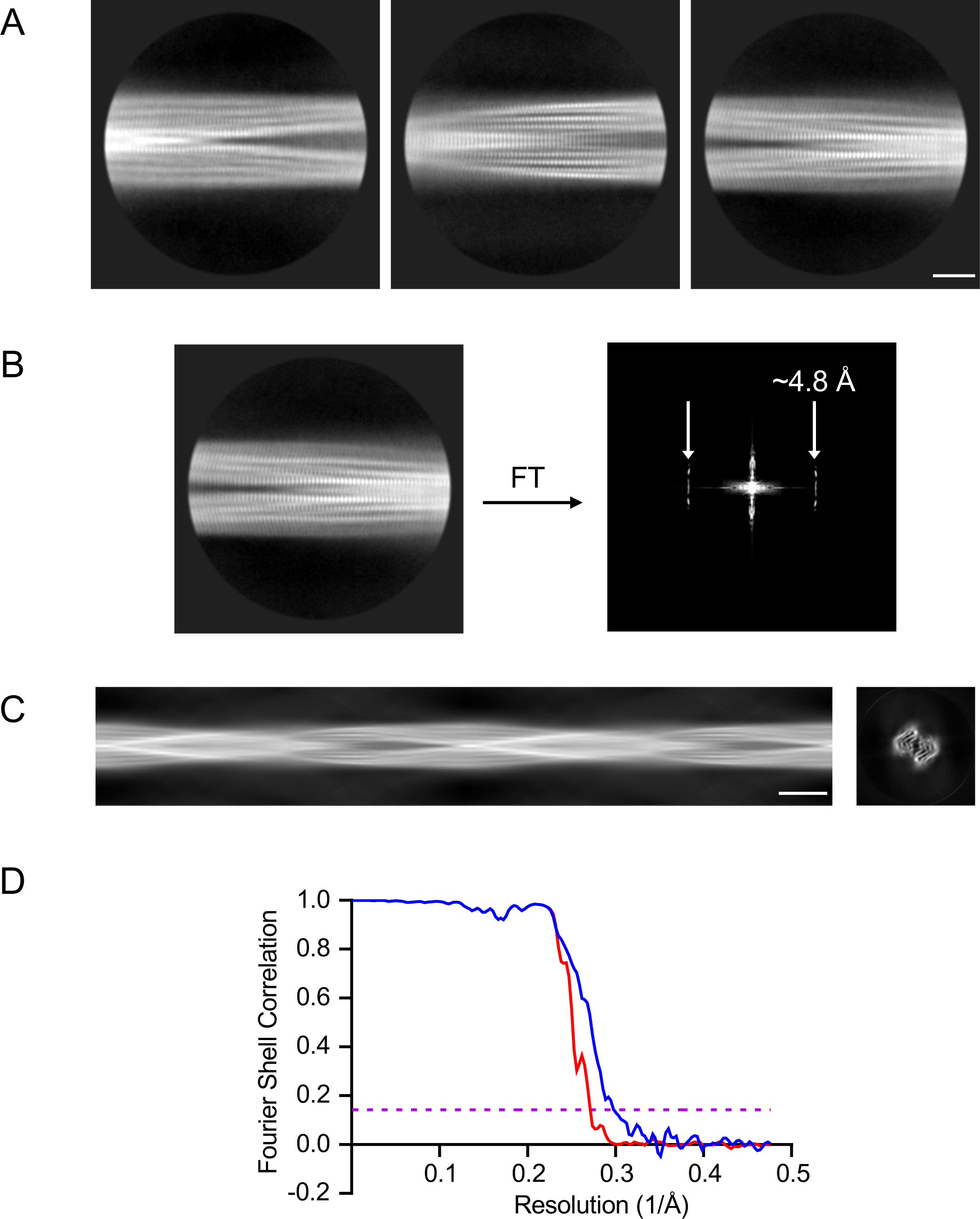
2D classes, cross-section and evaluation of the resolution of the cryo-EM high-resolution map of αSyn amyloid fibres in complex with WT DNAJB1. A) 2D classes of αSyn amyloid fibres in complex with WT DNAJB1. The DNAJB1 density was masked out to focus on fibre structure for this analysis. Scale bar, 50 Å. B) A 2D class and its corresponding FT, showing the 4.8 Å repeat. C) Side view of the aligned 2D classes and the calculated cross-section. Scale bar, 100 Å. D) Fourier shell correlation (FSC) curves of two independently refined half-maps with phase randomisation (red curve) and of the final cryo-EM reconstruction (blue curve). The dashed purple line indicates FSC = 0.143.

**Supplementary figure 4.**
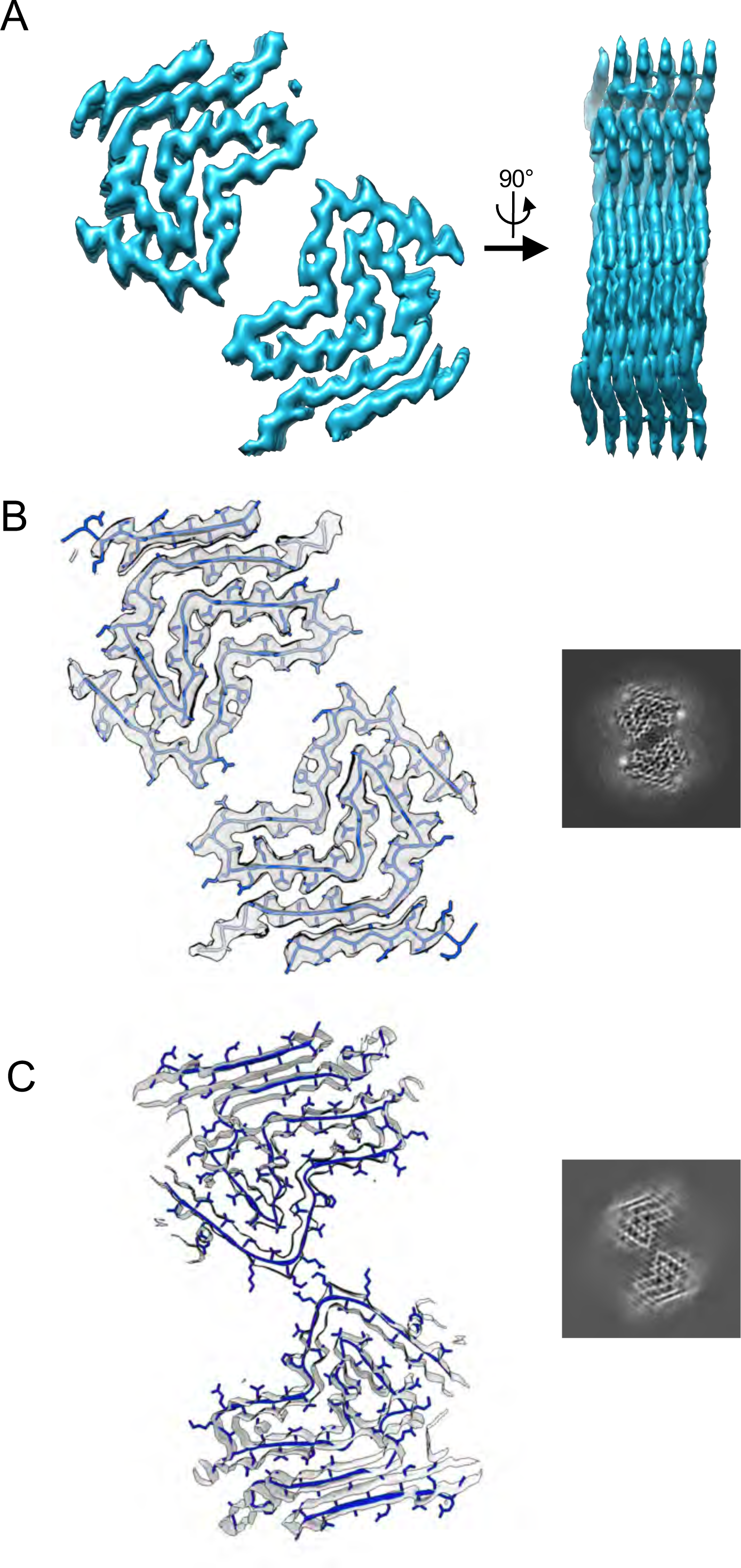
Structure of αSyn amyloid fibre from the WT DNAJB1 dataset. A) Cryo-EM map (cross-section and side view) of the αSyn amyloid fibre. B) Cryo-EM map of a single protofilament with the atomic model (PDB:6RT0) fitted into the density and the projected density slice of a single repeat.

**Supplementary figure 5.**
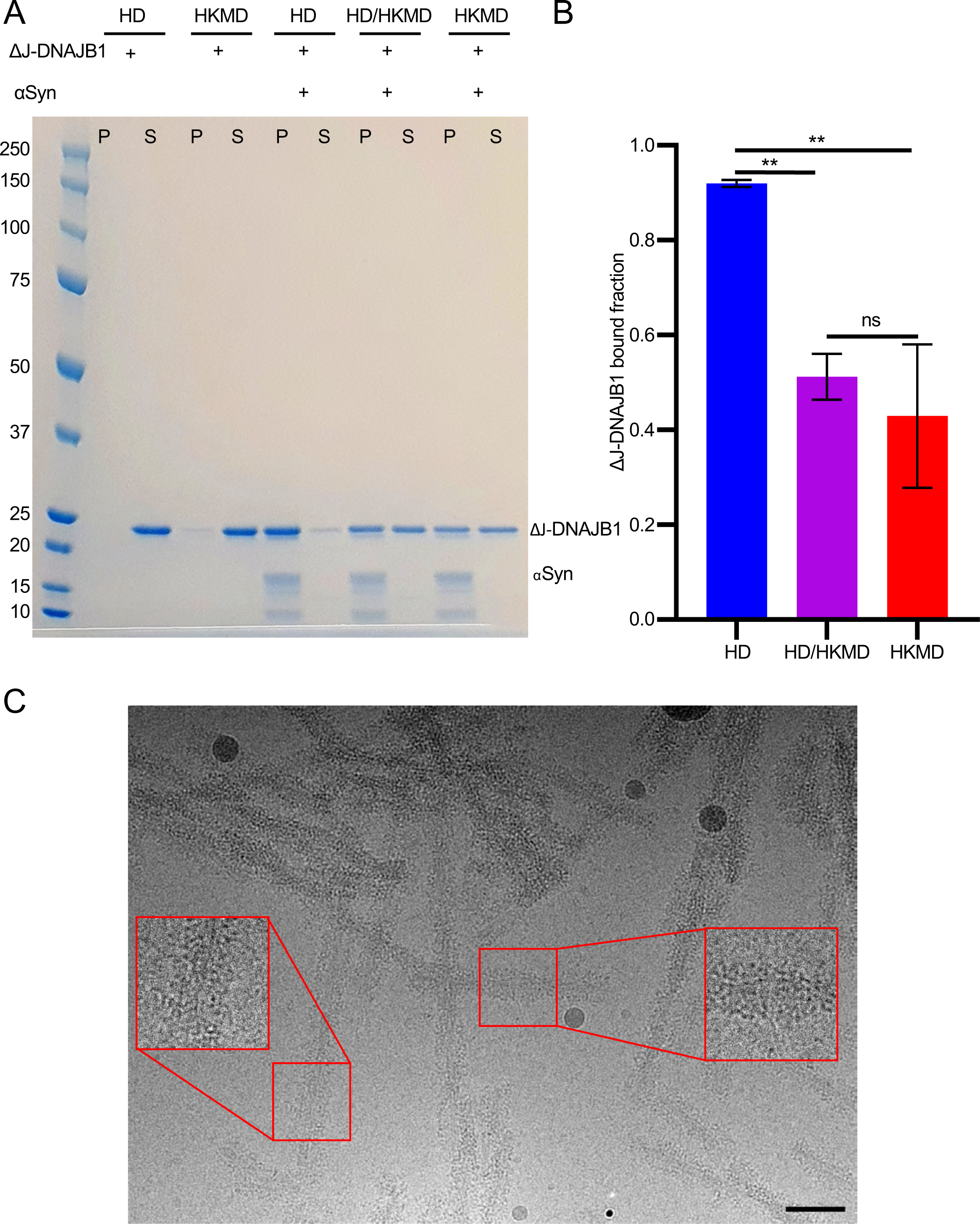
HD buffer also promotes ΔJ-DNAJB1 binding to αSyn amyloid fibres. A) Binding assay showing the reversibility of the binding when the salts are added back. The proteins were incubated twice in HD buffer (HD condition), twice in HKMD buffer (HKMD condition) or once in HD buffer and once in HKMD buffer (HD/HKMD buffer). B) Histogram of ΔJ-DNAJB1 bound fraction in each condition shown in (A) (N = 3 independent experiments). A Shapiro-Wilk test was performed to check the normality of the data, followed by a one-way ANOVA with Tukey’s multiple comparisons test (P = 0.0038 between HD and HD/HKMD conditions, P = 0.0015 between HD and HKMD conditions). C) Micrograph showing αSyn amyloid fibres decorated by ΔJ-DNAJB1 in HD buffer. Scale bar, 50 nm.

**Supplementary figure 6.**
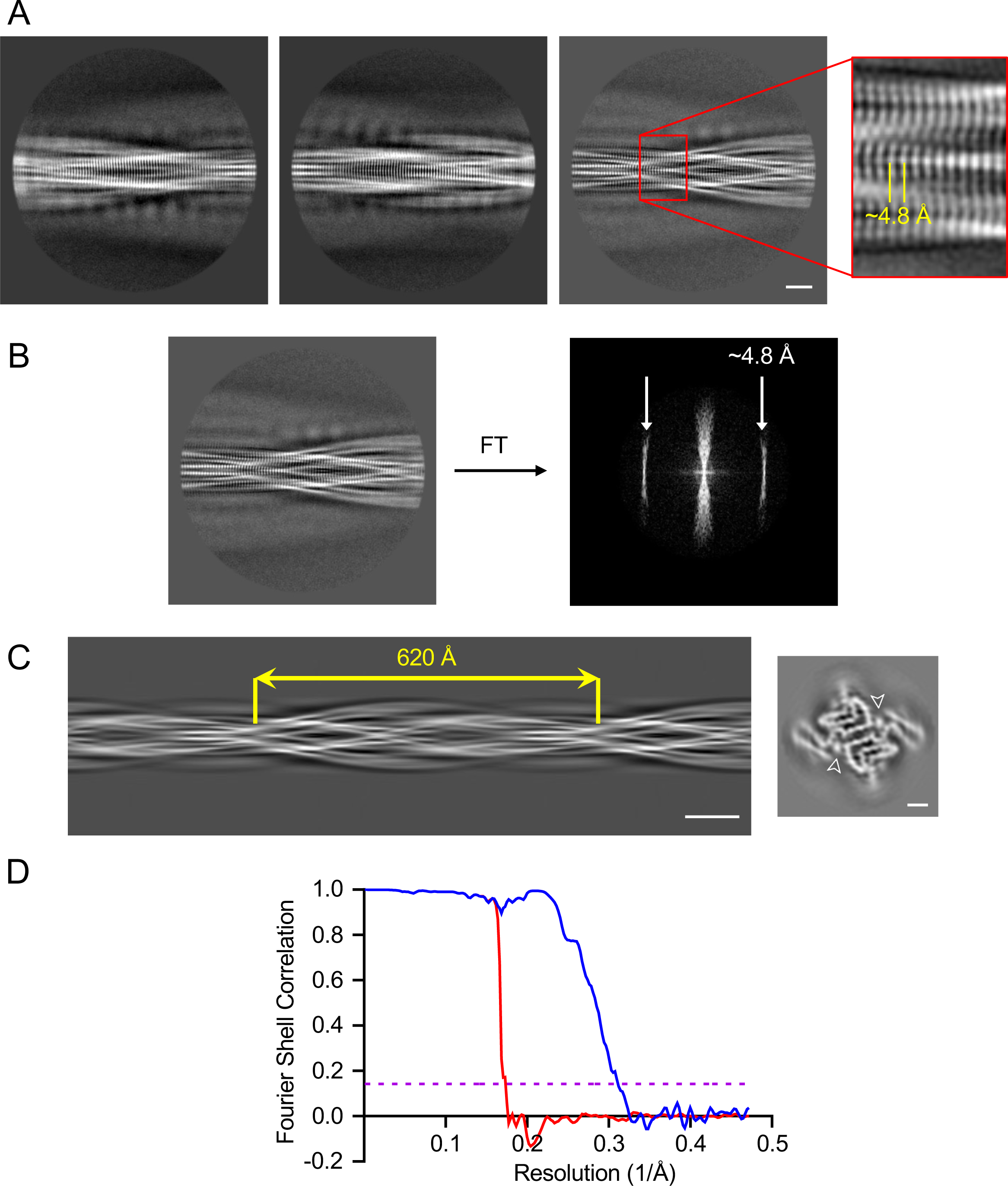
2D classes, cross-section and resolution plot of the cryo-EM map of αSyn amyloid fibres in complex with ΔJ-DNAJB1. A) 2D classes of αSyn amyloid fibres in complex with ΔJ-DNAJB1. A mask was applied to exclude ΔJ-DNAJB1 signal in the alignment. Scale bar, 50 Å. B) A 2D class and its corresponding FT, showing a peak at around 4.8 Å. C) Side view of the aligned 2D classes and the calculated cross-section. The crossover distance was estimated at 620 Å. Scale bars, 100 Å for the side view and 20 Å for the cross-section. D) Fourier shell correlation (FSC) curves of two independently refined, masked half-maps using phase randomisation (red curve) and of the final cryo-EM reconstruction (blue curve). The dashed purple line indicates FSC = 0.143.

**Supplementary figure 7.**
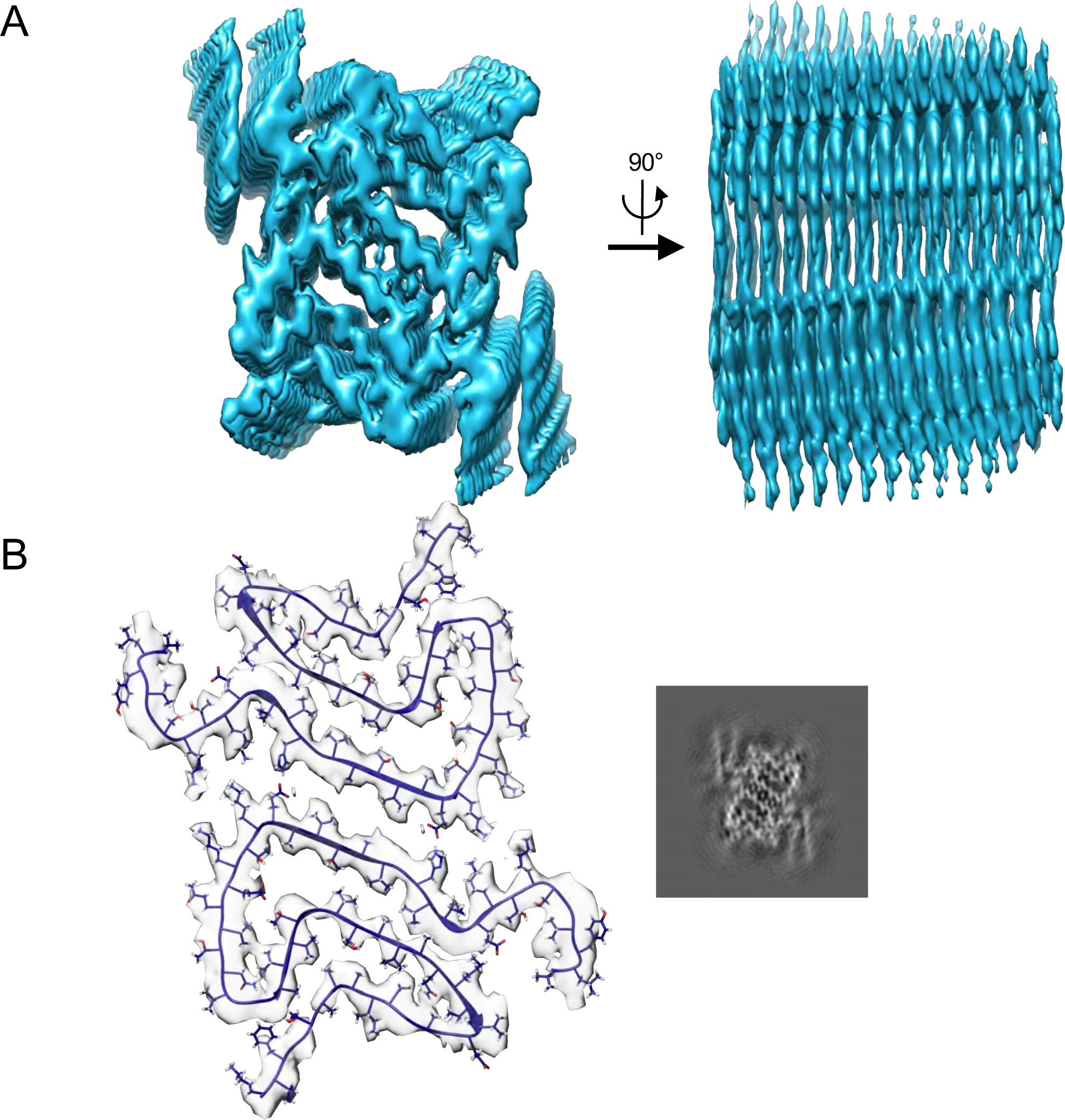
Structure of αSyn amyloid fibre from the ΔJ-DNAJB1 dataset. A) Cryo-EM map (cross-section and side view) of the αSyn amyloid fibre. B) Cryo-EM map of a single protofilament with the atomic model (PDB entry: 6OSJ) fitted into the density and the projected density slice of a single repeat.

**Supplementary figure 8.**
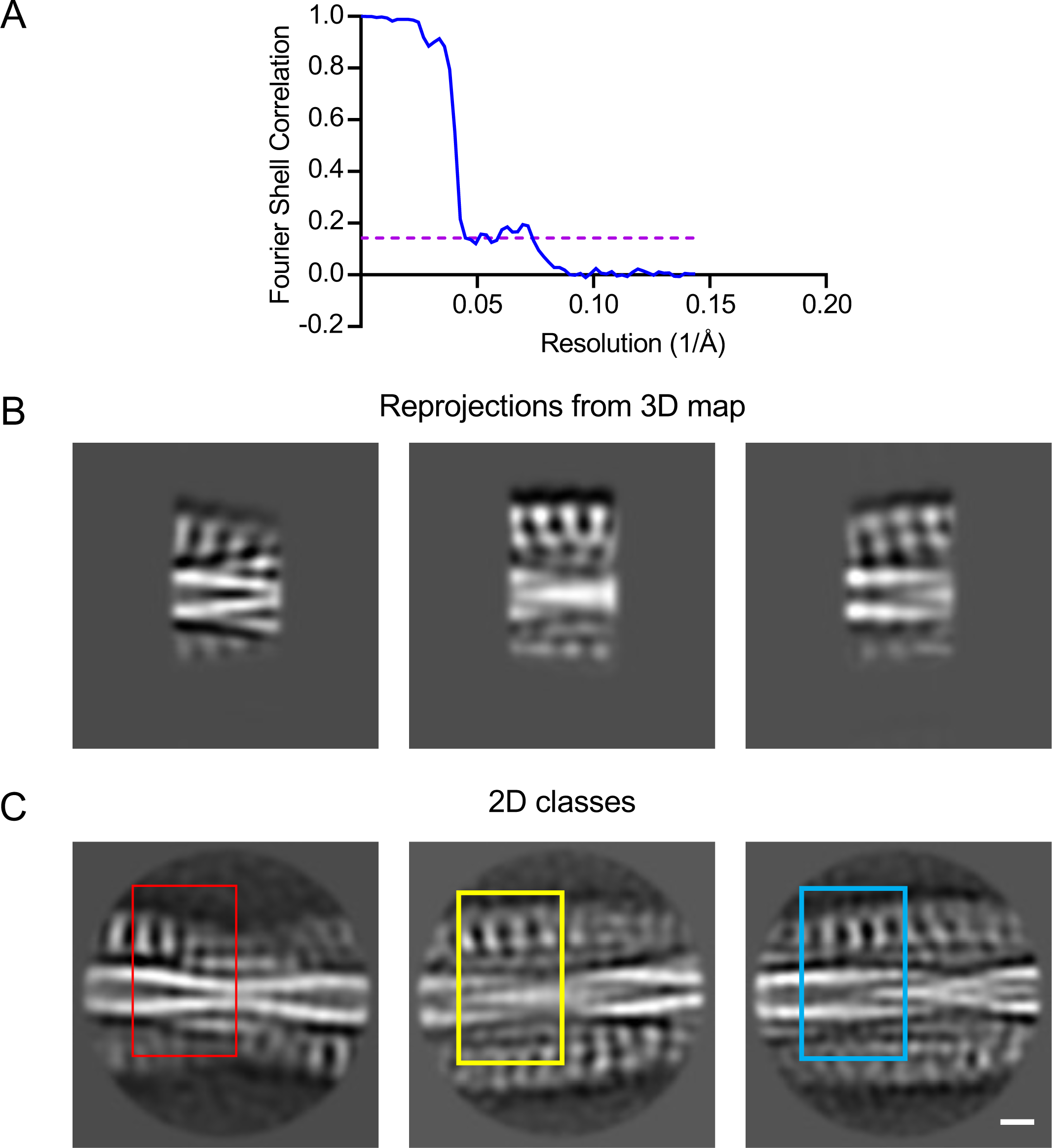
2D classes cross-section and resolution plot of ΔJ DNAJB1:αSyn fibre complex, 2D classes and reprojections from the map. A) FSC curve of the final cryo-EM reconstruction (blue curve). The dashed purple line indicates FSC = 0.143. B) Side-view reprojections from the postprocessed map. C) 2D class averages of the ΔJ DNAJB1:αSyn fibre complex. The 2D class averages resemble the reprojections. Scale bar, 50 Å.

**Supplementary figure 9.**
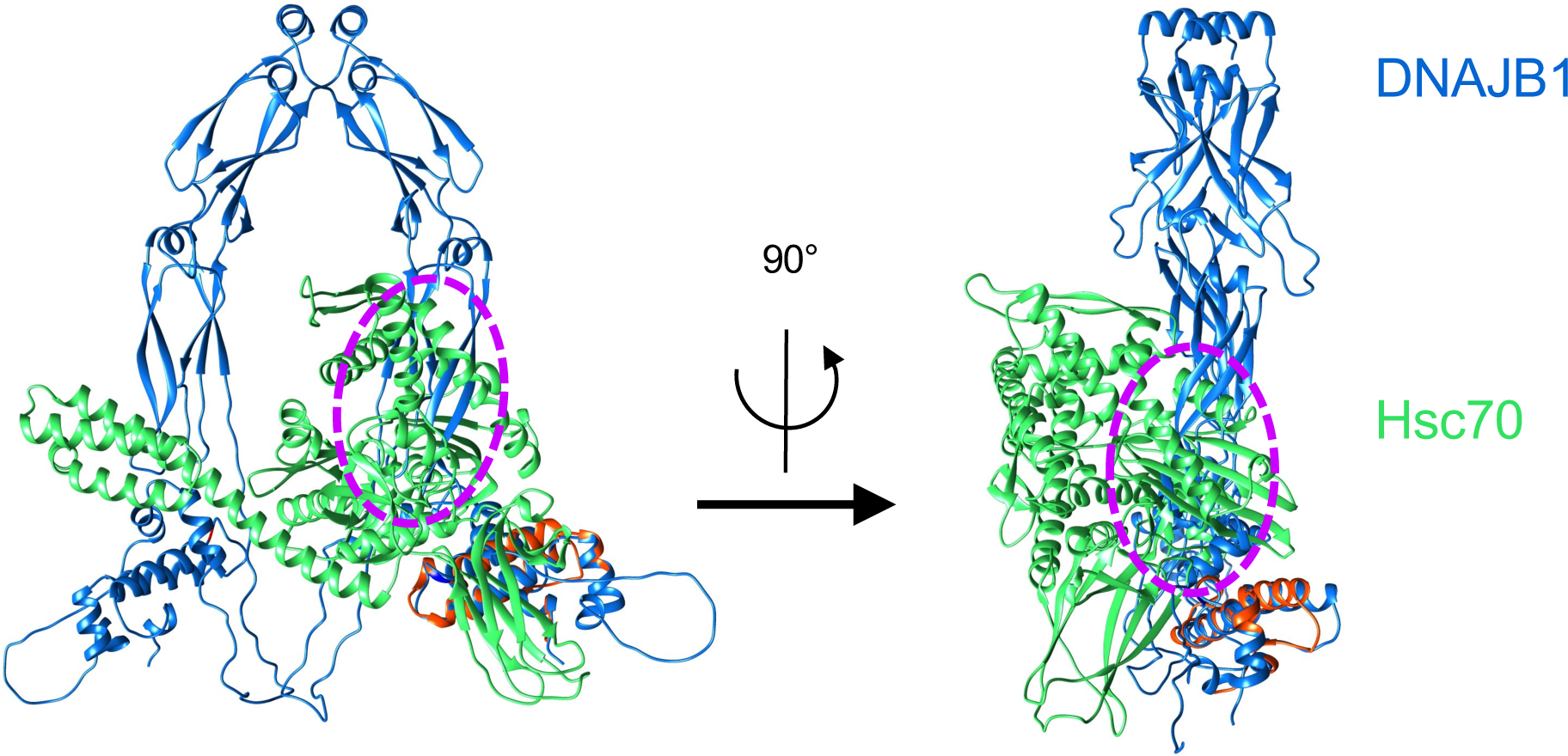
Clash between Hsp70 (green) and DNAJB1 in auto inhibited form (blue). They are aligned via the J domain present in both structures (red), using a DnaK-J domain structure (PDB 5nro) and the DNAJB1 fit from Figure 2. With the J domain close to the C terminal domain, the ATPase domain of Hsc70 totally overlaps with the C terminal domain (magenta dashed outlines).

